# Human Satellite 3 DNA encodes megabase-scale transcription factor binding platforms

**DOI:** 10.1101/2024.10.22.616524

**Authors:** J. Matthew Franklin, Danilo Dubocanin, Cy Chittenden, Ashlie Barillas, Rosa Jooyoung Lee, Rajarshi P. Ghosh, Jennifer L. Gerton, Kun-Liang Guan, Nicolas Altemose

## Abstract

Eukaryotic genomes frequently contain large arrays of tandem repeats, called satellite DNA. While some satellite DNAs participate in centromere function, others do not. For example, Human Satellite 3 (HSat3) forms the largest satellite DNA arrays in the human genome, but these multi-megabase regions were almost fully excluded from genome assemblies until recently, and their potential functions remain understudied and largely unknown. To address this, we performed a systematic screen for HSat3 binding proteins. Our work revealed that HSat3 contains millions of copies of transcription factor (TF) motifs bound by over a dozen TFs from various signaling pathways, including the growth-regulating transcription effector family TEAD1-4 from the Hippo pathway. Imaging experiments show that TEAD recruits the co-activator YAP to HSat3 regions in a cell-state specific manner. Using synthetic reporter assays, targeted repression of HSat3, inducible degradation of YAP, and super-resolution microscopy, we show that HSat3 arrays can localize YAP/TEAD inside the nucleolus, enhancing RNA Polymerase I activity. Beyond discovering a direct relationship between the Hippo pathway and ribosomal DNA regulation, this work demonstrates that satellite DNA can encode multiple transcription factor binding motifs, defining an important functional role for these enormous genomic elements.

## INTRODUCTION

Eukaryotic genomes often contain massive arrays of rapidly evolving repetitive DNA sequences called satellite DNA. The repeat units can range from a few bases to kilobases, and array lengths reach up to tens of megabases^1^. Although satellite DNAs were among the first sequences to be biochemically isolated and characterized in the early days of molecular biology, modern genomics research has largely excluded them^1^, and our understanding of the origins and evolutionary mechanisms that maintain these megabase-scale structures remains incomplete.

Despite the general notion that satellite DNAs are transcriptionally silent and constitutively heterochromatic, examples of transcription factors (TFs) binding satellite DNA through motif recognition have been reported for mouse cells (Pax/9, Foxd3, YY11^2^), human cells (HSF1^3^, DUX4^4^, Ikaros^5^, EWS-FLI1^6^), and *Drosophila*^7^. The biological functions of these interactions appear disparate, suggesting that the sequence content, chromatin composition, and genomic localization of satellite DNAs dictate the functional roles of these massive repetitive elements.

Recent efforts using long-read sequencing have established complete “Telomere-to-Telomere” (T2T) assemblies of humans^8^ and non-human primates^9^. These T2T assemblies provide unprecedented characterizations of satellite DNAs in terms of sequence content and variation between individuals. In humans, satellite DNAs are predominantly found at centromeres, peri-centromeres, the short arms of the acrocentric chromosomes (13-15, 21-22), and the long arm of chromosome Y. In total, this accounts for over 200 megabases, approximately 7% of the human genome^10^.

To date, satellite DNA studies in humans traditionally focused on alpha satellite (αSat), the most abundant satellite DNA family, which provides an essential scaffold for assembly of the centromere and kinetochore on each chromosome. Recently, a search for αSat binding transcription factors revealed multiple new interactions, including BRD4, which normally functions at enhancer elements^11^. The other major human satellite DNA families (Human Satellites 1, 2, 3, beta, gamma) have poorly understood biological functions, and their heterogeneous localization across chromosomes suggests they are not essential for centromere function.

Human Satellite 3 (HSat3) is the most abundant non-centromeric satellite in the human genome, representing approximately 2% of the genome. This family is defined by enrichment of GGAAT pentamers and is thought to have expanded along the great ape lineage^1^. The largest contiguous repetitive element in the human genome is an HSat3 array adjacent to the chromosome 9 centromere, spanning 28 megabases^10^. Notably, the acrocentric chromosomes (13-15, 21, 22) contain large arrays of ribosomal RNA genes (rDNAs) that are bookended by all the major satellite DNA repeat families^10^, including dozens of distinct HSat3 arrays ranging from ∼1 kilobase to 8 megabases (Supplementary Figure 1). HSat3 DNA is generally thought to be packaged as constitutive heterochromatin, marked by high levels of methyl-CpG and H3K9me3-containing nucleosomes, and maintained in a silenced, compacted state^12^. However, a handful of reports describe enrichment of transcription factors (TFs) on HSat3^3,6,13,14^.

Here, we report the results of a systematic search for HSat3-binding TFs using motif enrichment analysis and published ChIP-seq data. This effort revealed a set of HSat3-binding TFs from multiple, broadly important cell signaling pathways. We conducted functional studies of the top hit, TEAD, a highly conserved TF family from the Hippo pathway that plays vital roles in early development, organ size regulation, and cancer^15,16^. Based on evidence from super-resolution microscopy, RNA Polymerase I reporter assays, and genetic perturbations, we propose that HSat3-bound TEAD and its effectors regulate rDNA transcription in the nucleolus.

## RESULTS

### A transcription factor screen reveals candidate HSat3 binding factors

To systematically identify novel HSat3-binding transcription factors, we first scanned HSat3 arrays for known transcription factor motifs from the JASPAR database (828 vertebrate motifs)^17^. Annotated HSat3 sequences were sourced from the human T2T assembly T2T-CHM13v2.0^1^ (Figure 1a). Although closely related, distinct HSat3 arrays within humans show sequence divergence. Using k-mer based clustering, HSat3 repeats can be split into two main subfamilies, HSat3A and HSat3B, each containing further subfamilies (HSat3A1-6, HSat3B1-5)^18^. We computed the enrichment of each known TF on each distinct HSat3 array, relative to the motif’s frequency in gene promoters (Figure 1b). As expected, most known TF binding motifs are not enriched, given the highly constrained sequence space of repeat elements (Figure 1c, Supplementary data). However, we found that 42 TF motifs are enriched at least 10-fold compared to background, sometimes with HSat3 subfamily-specific enrichment profiles (Figure 1c).

**Figure 1:**
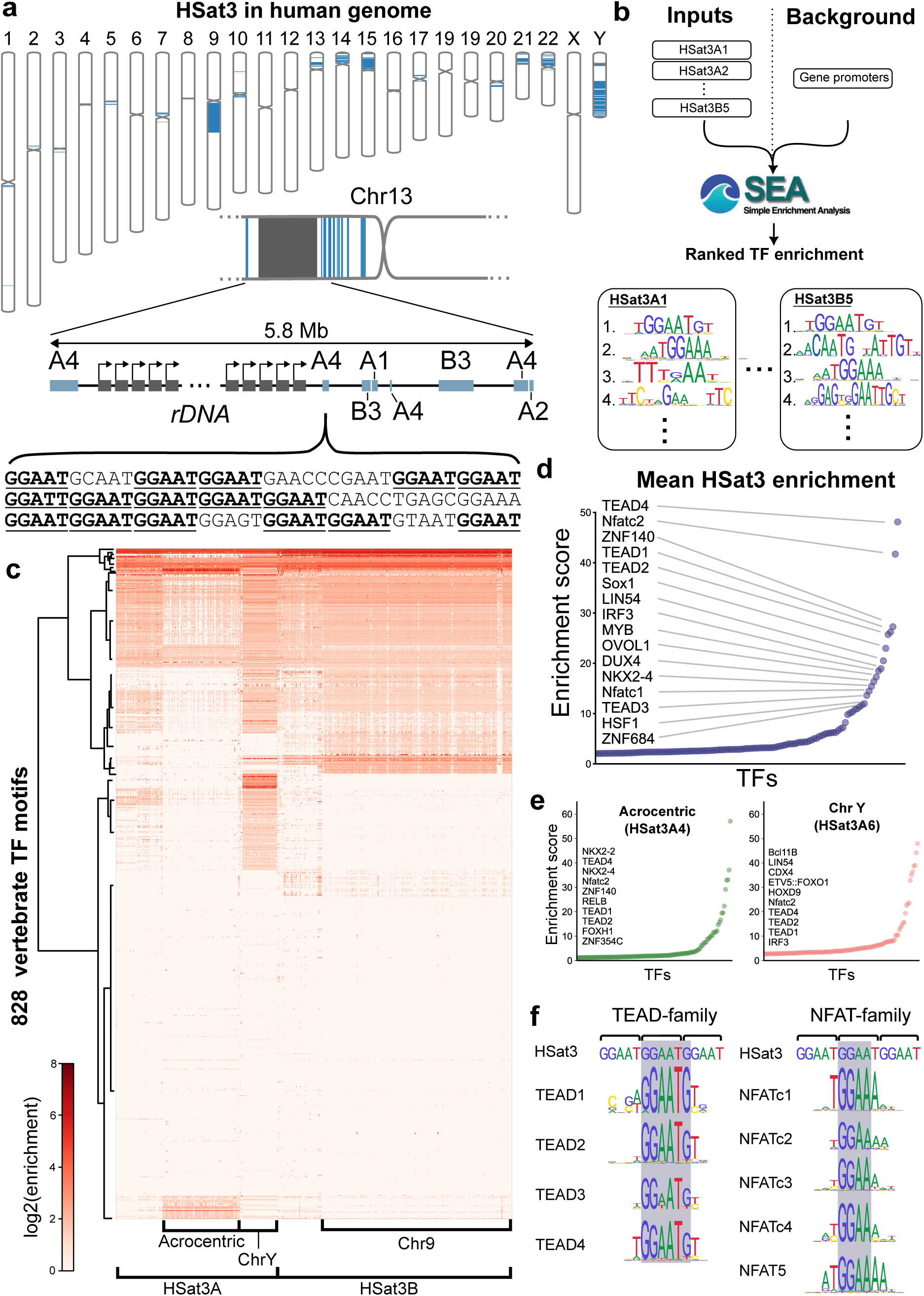
A search for transcription factor motifs enriched in HSat3. (a) Top: karyogram of the human genome with HSat3 arrays marked in blue. Note the large arrays on chromosomes 9, 15, Y. Middle: Depiction of HSat3 (blue) and rDNA (grey) organization on chromosome 13 p-arm. Individual HSat3 arrays are annotated with subfamily names. Bottom: sample of HSat3A4 DNA sequence with GGAAT pentamers in bold with underline. (b) Flow diagram of the motif enrichment search using SEA from MEME-Suite. (c) Heat-map of predicted TF motifs enriched on each HSat3 array instance in the chm13v2.0 genome. Columns are individual HSat3 arrays, rows are individual TF motifs organized by hierarchical clustering. (d) Mean enrichment scores for each TF across all HSat3 arrays, with labels for the top 16 TFs. (e) HSat3 sub-family specific enrichment scores with top 10 motifs labeled for the acrocentric-specific subfamily HSat3A4 and the chromosome Y-specific subfamily HSat3A6. (f) Correspondence between the HSat3 pentamer repeat and the TEAD- and NFAT-family consensus motifs (logo diagrams from JASPAR^17^).

Across all HSat3 arrays, the TEAD and NFAT TF families represent the most highly enriched motifs, followed by SOX1, LIN54, IRF3, MYB, OVOL1, DUX4, and NKX2-4 (Figure 1d). Previously described as an HSat2 binding factor, the DUX4 motif is also highly enriched in HSat3. Both HSat2 and HSat3 are enriched for GGAAT pentamers, but HSat2 has a longer, variable repeat unit which has diverged from HSat3^19^. Although previously validated^20^, HSF1 is ranked 15th in terms of motif enrichment. Although the top enriched motifs are generally consistent among HSat3 subfamilies (Figure 1e, Supplementary Figure 2), the Y-chromosome specific subfamily HSat3A6 contained unique motif enrichments for bcl11B, CDX4, ETV5-FOX01, and HOXD9 (Figure 1e). This is consistent with the strong sequence divergence of HSat3A6^1^. Surprisingly, ETV5-FOX01 is the only ETS-family motif heavily enriched in HSat3, despite the core GGAA-motif found in most ETS consensus motifs^17^ (Supplementary Figure 3).

Notably, the well-characterized TEAD DNA binding motifs contain a constrained core GGAATG, which is recapitulated by tandemly repeated GGAAT HSat3 pentamers (Figure 1f). Similarly, NFAT-family motifs consistently contain GGAA in the consensus motif (Figure 1f). We estimated that the percentage of total TEAD motifs across the human genome in HSat3 ranges from 24-58% using ChIP-seq derived motifs in JASPAR (Supplementary Figure 4). However, all TEAD-family factors contain a highly constrained GGAATG 6 base pair core. Approximately 72% of genome-wide matches to these core motifs occur in HSat3 (Supplementary Figure 4).

As many of these candidate factors are not known to bind HSat3, we turned to existing ChIP-seq data to corroborate our predictions. Standard ChIP-seq pipelines mask out repetitive regions, possibly explaining the dearth of established HSat3 factors despite the strong enrichment of TF motifs. The ENCODE database has standardized high-quality ChIP-seq datasets across many cell types and factors^21^. To utilize this rich dataset we re-mapped the data to theT2T-CHM13v2.0 assembly^22^. A similar approach was previously taken to identify factors that bind αSat^11^.

We defined the enrichment score as the coverage ratio of HSat3 arrays to random control regions for the ChIP pull-down and the input DNA (Figure 2a, Methods, Supplementary Data). This controls for specificity in both the ChIP pull-down and the input DNA. Mapping short-read sequencing data to repetitive DNA is inherently limited due to multiple mapping sites. To examine our ability to map short reads within HSat3 arrays, we used existing Heat-shock Factor 1 (HSF1) ChIP-seq data from heat-shocked MCF7 cells^23^. HSF1 is well described as an HSat3-binding TF under cellular stress conditions, most notably following heat-shock^3^. Interestingly, HSF1 motifs were enriched in sub-regions of HSat3 arrays, in contrast to TEAD motifs, which are evenly distributed (Figure 2b). The HSF1 ChIP-seq data showed clear enrichment at HSat3 regions of high motif density. This example shows that short-read sequencing data can map broadly to predicted regions even within simple repeat arrays.

**Figure 2:**
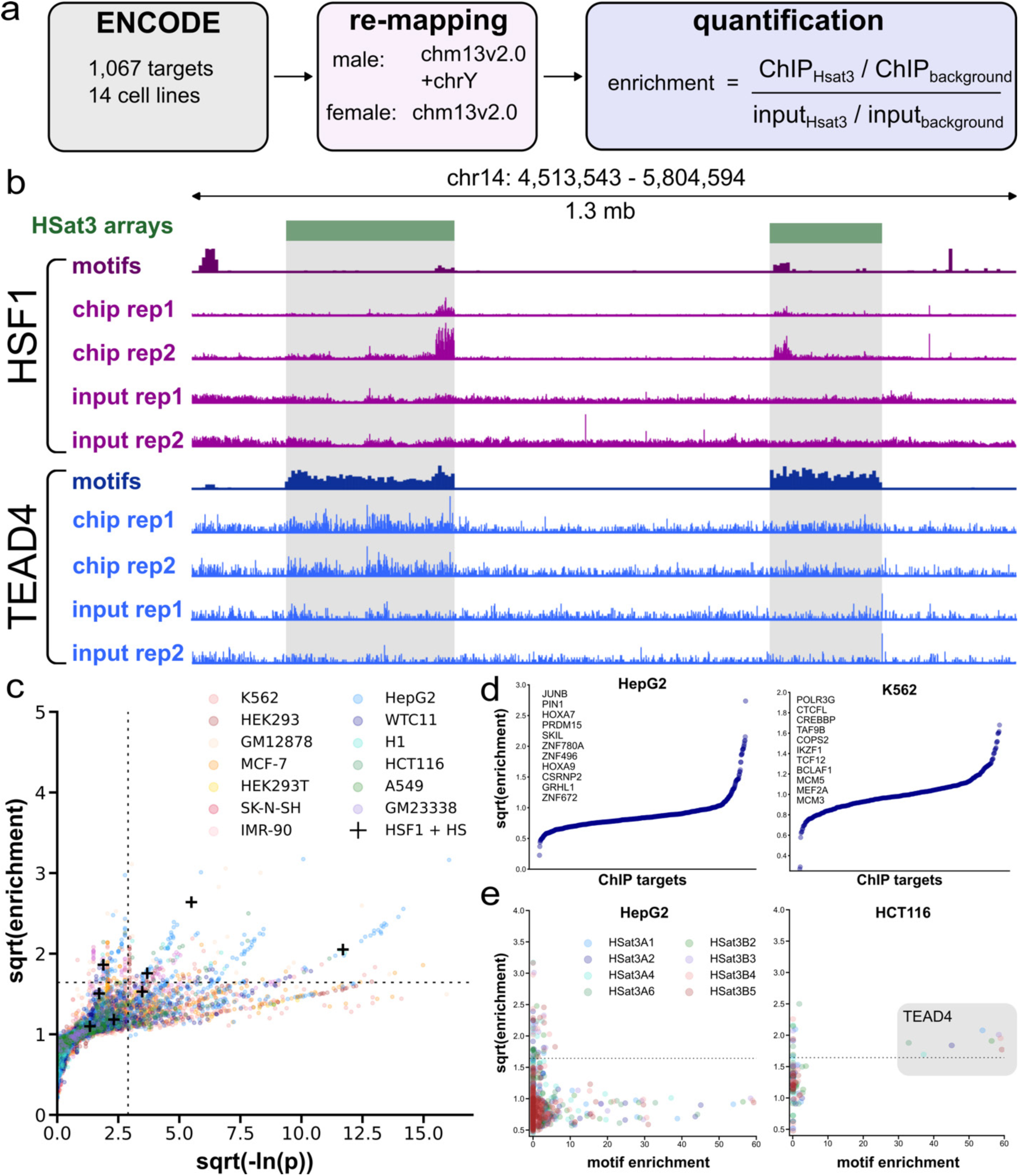
ENCODE ChIP-seq database search for HSat3-binding factors. (a) Flow diagram of ChIP-seq analysis pipeline. Data were accessed directly from ENCODE. 1,067 ChIP-targets across 13 human cell lines were mapped to the T2T-CHM13v2.0 assembly with or without chromosome Y depending on cell line karyotype. Enrichment on HSat3 was calculated using the ratio of HSat3 to background for the ChIP relative to the input DNA. (b) Demonstration of ChIP-seq reads enriched at TF motifs in HSat3. HSF1 motifs are found in sub-regions of the HSat3 arrays (green blocks). HSF1 ChIP-seq reads in heat-shocked MCF7 cells show specific enrichment in regions of high HSF1 motif density. TEAD4 motifs are enriched evenly across HSat3 arrays, and TEAD4 ChIP-seq in HTC116 cells shows read enrichment uniformly across HSat3. (c) Scatterplot of enrichment vs -ln(P) for the ChIP-seq screen. Data points are colored by cell line with markers for each HSat3 subfamily. ChIP-seq for HSF1 after heat-shock in MCF7 cells is highlighted with “+” markers as a positive control. (d) Ranked HSat3 enrichment scores for HepG2 (668 targets) and K562 (468 targets). (e) Correlation between ChIP-seq and motif enrichment for HepG2 (N=266 factors) and HCT116 (N=21 factors), colored by HSat3 subfamily. Data are plotted for subfamilies with more than 10 instances in the reference genome.

The enrichment scores for 1,067 targets across 13 commonly used cell lines revealed dozens of highly enriched factors (Figure 2c-d, supplementary data). The top 10 factors are mostly cell-line specific (Supplementary Figure 5), possibly reflecting cell-type specific functionality and TF expression.

Across cell lines, the only target showing HSat3 enrichment in both ChIP-seq and motif analysis was TEAD4 in HCT116 (Figure 2e, Supplementary Figure 6). This highlights the complementarity of the two approaches and some of the limitations of screens based on ChIP-seq enrichment in aggregate. For example, NFAT family members, which were highly enriched in the motif enrichment screen, were not found among the top TF candidates for any cell lines in our ChIP-seq screen. While TEADs are broadly expressed in most cell lines and constitutively nuclear, NFAT factors are only translocated into the nucleus following intracellular calcium activation via calcineurin^24^. The NFAT ENCODE datasets were not carried out in calcium activating conditions, thus potentially explaining the lack of HSat3 signal. We next sought to validate the top HSat3 candidates from both the motif and ChIP-seq analyses.

### Imaging and sequencing validate TFs bound to HSat3

Immunostaining for HSF1 after heat shock shows distinct sub-nuclear foci due to the high density of binding sites on large HSat3 arrays^3^. Similarly, live imaging of dCas9 targeted to the chromosome 9 HSat3 array shows clear foci^25^. Thus, we reasoned that we could leverage the ability to image HSat3-associated foci to test our binding predictions. To that end, we cloned a library of the top candidate TFs tagged with mScarlet for transfection and live-cell imaging in 293T cells. Further, we generated a synthetic zinc-finger (ZF) array targeting 18 bp of the repeated HSat3 pentamer GGAAT to provide a colocalization fiducial, referred to as ZFsat3. ZFsat3-mScarlet shows punctate foci in live cells, and two-color immuno-FISH with pre-labeled anti-RFP-CF488A antibody and fluorophore-labeled HSat3 oligos showed a one-to-one colocalization, demonstrating highly specific HSat3 binding (Supplementary Figure 7).

Each mScarlet-tagged candidate TF was transiently transfected with ZFsat3-meGFP to assess HSat3 localization using live-cell microscopy (Figure 3a). As a positive control, we show previously identified HSat3 factors JRK and TIGD3 both co-localize with HSat3 foci (Supplementary Figure 8). TEAD1-mScarlet showed a one-to-one correspondence to ZFsat3 (Figure 3b), consistent with the near-perfect recapitulation of the TEAD consensus motif by HSat3 pentamers (Figure 1f). Similarly, TEAD1-4 tagged with meGFP showed similarly strong colocalization with ZFsat3-mScarlet (Supplementary Figure 8). TEAD-bound sites frequently contain multiple TEAD motifs in human enhancers^26^, with the most common configuration being two motifs separated by 3 base pairs. Cooperative TEAD binding is suggested to occur at such repeated motifs, thus accumulation of TEADs at HSat3 may be driven by cooperativity.

**Figure 3:**
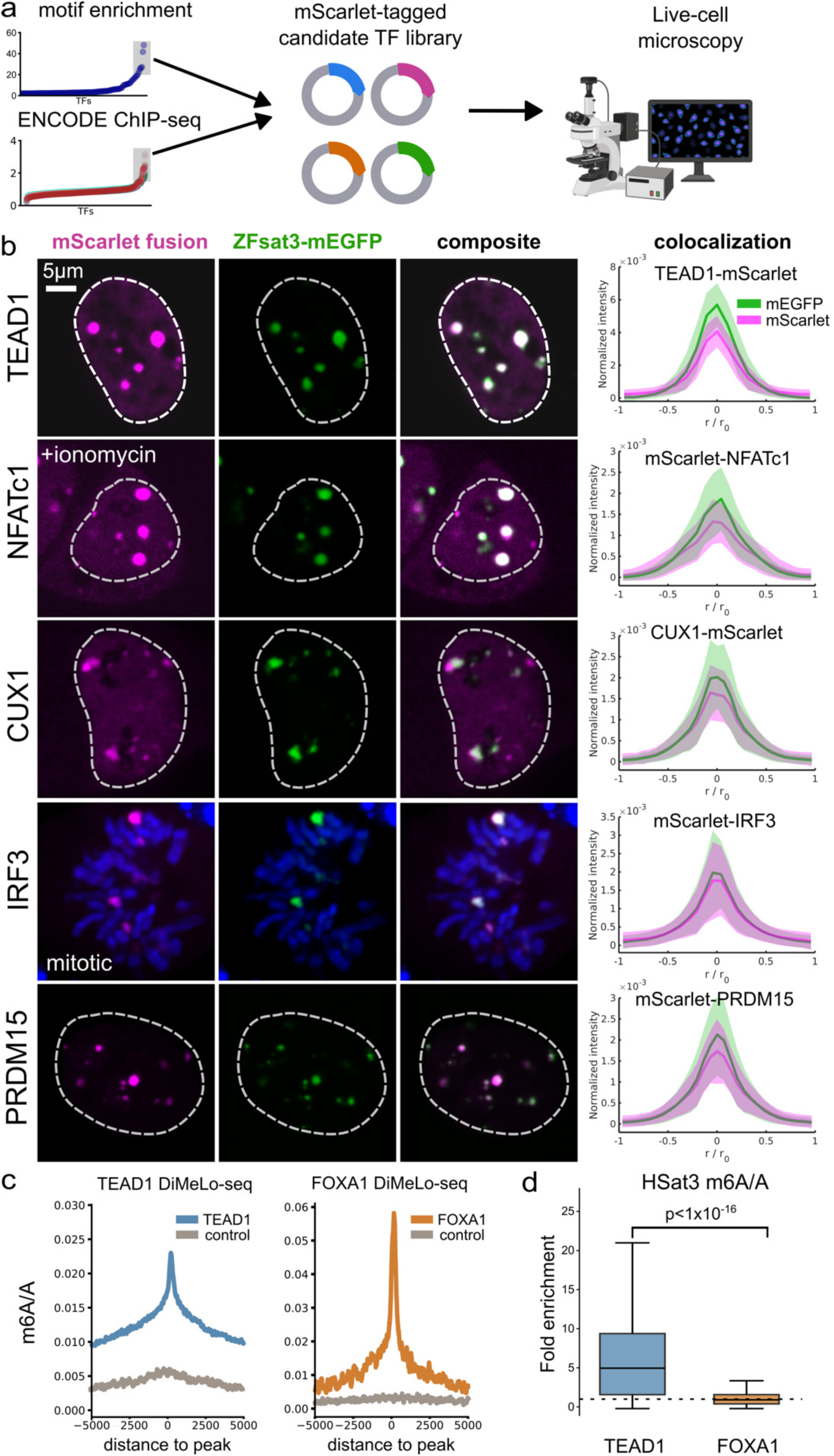
Validation of predicted HSat3-binding factors. (a) Flow diagram of the validation pipeline. Top candidate factors from both motif searches and ChIP-seq analysis were tagged with mScarlet in a pHIV EF1ɑ lentivirus expression vector. Transfected candidate TFs were evaluated with live-cell microscopy. (b) Representative confocal images of TF-mScarlet and ZFsat3-mEGFP co-transfected in 293T cells for validated HSat3 factors TEAD1, NFATc1, CUX1, IRF3, and PRDM15. All images are single z-planes. Due to the high binding capacity for HSat3, foci intensities are extremely bright compared to nucleoplasm, thus outlines of cell nuclei are shown by dashed lines. Right panels show intensity profile analysis of 3D segmented HSat3 foci with normalized intensity and the spatial dimension. The number of foci used for each distribution are as follows: N(TEAD1)=10, N(NFATc1)=20, N(CUX1)=15, N(IRF3)=12, N(PRDM15)=15. The solid line represents the mean and the shaded regions represent the standard deviation. (c) Results of DiMeLo-seq for TEAD1-mScarlet (HSat3 binding), FOXA1-mScarlet (non-HSat3 targeting), and wildtype 293Ts not expressing mScarlet. The fraction of m6A-modified bases (m6A/A) was calculated for reads centered at the respective ChIP-seq peaks for TEAD1 and FOXA1 in non-repetitive regions. (d) Enrichment of m6A/A across all HSat3-aligned DiMeLo-seq reads for either TEAD1 and FOXA1, relative to control cells. The box plot lines are as follows: center line, median; box limits, upper and lower quartiles; whiskers, 1.5x interquartile range. A one-tailed Wilcoxon rank sum test was performed to compare m6A/A enrichment in HSat3 between TEAD1 and FOXA1.

NFAT factors are cytoplasmically localized until activation by intracellular calcium release^24^. Following 1 hr treatment with 1 μM ionomycin, mScarlet-tagged NFATc1-3 and NFAT5 showed nuclear translocation in cells to HSat3 foci (Figure 3b, Supplementary Figure 9a). In the Protein Atlas database, NFATc1 and NFATc4 both show clear nuclear bodies^27^. Although NFAT localization to such bodies has long been observed in fixed and live cell images^28^, our data provide an underlying molecular explanation.

Following activation of the core innate immunity response pathway, IRF3 is translocated into the nucleus to drive interferon expression^29^. Live imaging of mScarlet-IRF3 showed cytoplasmic localization in interphase 293T cells (Supplementary Figure 9b). However, mitotic cells showed strong IRF3 enrichment at HSat3 foci (Figure 3b). Sub-nuclear IRF3 foci have previously been identified and attributed to liquid-liquid phase separation with interferon (IFN)-stimulated response element DNA^30^. Upon inspection, the previously observed IRF3 foci resemble HSat3 foci, with tens of nuclear foci of variable size. Our results suggest that HSat3 provides a scaffold of dense binding motifs that drive this observed formation of nuclear foci.

The other evaluated top candidates SOX1 and NKX2-2 did not show overt localization to all ZFsat3-foci, however both SOX1 and NKX2-2 showed colocalization to a small subset of ZFsat3 foci (Supplementary Figure 10). Surprisingly, the CUT homeo-domain factor CUX1 showed strong colocalization with ZFsat3 (Figure 3b), despite being ranked 31^st^ for mean HSat3 binding in the motif-enrichment screen. The other CUT family members CUX2, ONECUT1, ONECUT2 have similar DNA binding domains and motifs, suggesting this family is likely to bind HSat3 (Supplementary Figure 11). Although the ZFsat3-megfp may be competing with HSat3-binding TFs, our results highlight that simple motif enrichment predicts the binding of many satellite DNA TFs.

Because the ChIP-seq screen revealed tens of candidate factors per cell line, we filtered our predictions using microscopy images showing protein localization to nuclear bodies or foci from Protein Atlas^27^ and existing literature. Ranked 4th in HepG2 (enrichment ratio=4.34), published PRDM15 immunofluorescence images show striking localization to nuclear bodies^27,31^. In our colocalization assay, PRDM15 showed strong colocalization to HSat3 (Figure 3b). The JASPAR PRDM15 motif does not enrich in HSat3 (rank= 751/868). However, a reported consensus motif from mouse ChIP-seq data shows highly constrained AA and GG dinucleotides separated by 5 bases^32^.

Flexibility in the ZF array may allow PRDM15 to bind AA and GG dinucleotides separated by 6 bases, which is found in to end-to-end HSat3 pentamers. We hypothesized that HSat3 localization may be mediated through the ZF array present on the C-terminus of PRDM family factors^33^. Truncated PRDM15 containing only the ZF array localized to HSat3, further suggesting direct binding via DNA motif recognition, although ZF-protein interactions cannot be ruled out (Supplementary Figure 12). Such flexible binding modes are reported for ZNF512 and ZNF512B, which have longer linkers between ZFs, aiding their ability to recognize repetitive DNA between species^34^.

Other tested ChIP-seq candidates JUNB, HOXA9, HOXA7, and HOXA9 did not show HSat3 co-localization in 293T transient transfection (Supplementary Figure 13). In the future, expanding the screen to test the ChIP-seq hits using the respective cell lines in which the factors showed HSat3 enrichment will be needed to thoroughly capture all HSat3-binding factors. Given the strong HSat3 colocalization in our imaging screen, the lack of NFAT and TEAD enrichment in ChIP-seq could be due to variations between cell lines, antibodies, and sample preparations.

We further validated TEAD binding to HSat3 using DiMeLo-seq. This technique maps protein-DNA interactions by recruiting an adenine DNA methyltransferase to a target protein, depositing exogenous methyl-6-adenine (m6A) marks, and reading the modified bases using long-read sequencing^35^. This allows accurate mapping of reads to repetitive regions of the genome. Anti-mScarlet DiMeLo-seq on TEAD1-mScarlet expressing 293T cells showed clear enrichment at known TEAD ChIP-seq peaks (Figure 3c), and a 6.25-fold increase in HSat3 methylation compared to control cells, whereas the non-HSat3 enriched TF FOXA1 showed only a 1.59-fold increase in HSat3 methylation (Figure 3d).

Our data show that repetitive DNA can act as an endogenous recruitment platform for many TFs, explaining previous observations of nuclear-body localized PRMD15 and NFAT-family factors. However, our catalog of HSat3-binding factors is not exhaustive. The motif enrichment strategy employed is relatively simple, and would not account for non-linear effects like binding cooperativity due to proximity of multiple weak sites. Further, these identified interactions imply that ChIP-seq derived motifs may be biased in predicting the true consensus sequence due to ignoring binding sites in repetitive regions.

Utilizing more datasets for protein-DNA interactions and proteomics-based approaches may reveal more candidate factors for HSat3 and other satellites. Because many nuclear factors are described as localizing to nuclear ‘speckles’ or ‘bodies’, it is likely that a subset of these observations are explained by massive repetitive arrays enriched for binding motifs.

The validated HSat3-binding factors are associated with a wide range of signaling pathways and disease states, with no obvious shared functionality. While TEADs are the effector of the Hippo pathway, NFAT members play important roles in T-cell activation and development^36^. IRF3 drives the interferon expression following innate immunity activation^37^. CUX1 is a broadly expressed repressor with specific roles in neuronal development^38^. PRDM15 maintains naive pluripotency via WNT / MAPK-ERK signaling pathways^32^. Thus, the precise functions of each potential TF-HSat3 interaction need to be evaluated separately. We focused our attention on our top hit, the TEAD family, given its well-studied roles in development, tissue homeostasis, and disease^39^.

### Endogenous TEAD1 binds HSat3

TF binding to HSat3 could be exaggerated by overexpression, given the high density of potential binding sites and the prior expectation that these regions are maintained in a dense heterochromatic state. We therefore generated MCF10A cells in which we tagged endogenous TEAD1 with eGFP to assess its localization to HSat3. Indeed, confocal imaging revealed multiple TEAD1 foci in each cell nucleus (Figure 4a). The largest foci likely correspond to the large HSat3 array on chromosome 9 (Figure 1a), as this XX line does not have the large chromosome Y array. To confirm endogenous TEAD1 foci are indeed binding HSat3, we generated metaphase spreads with MCF10A TEAD1-egfp cells, probing with 488-Alexa conjugated anti-GFP nanobody (Figure 4b).

**Figure 4:**
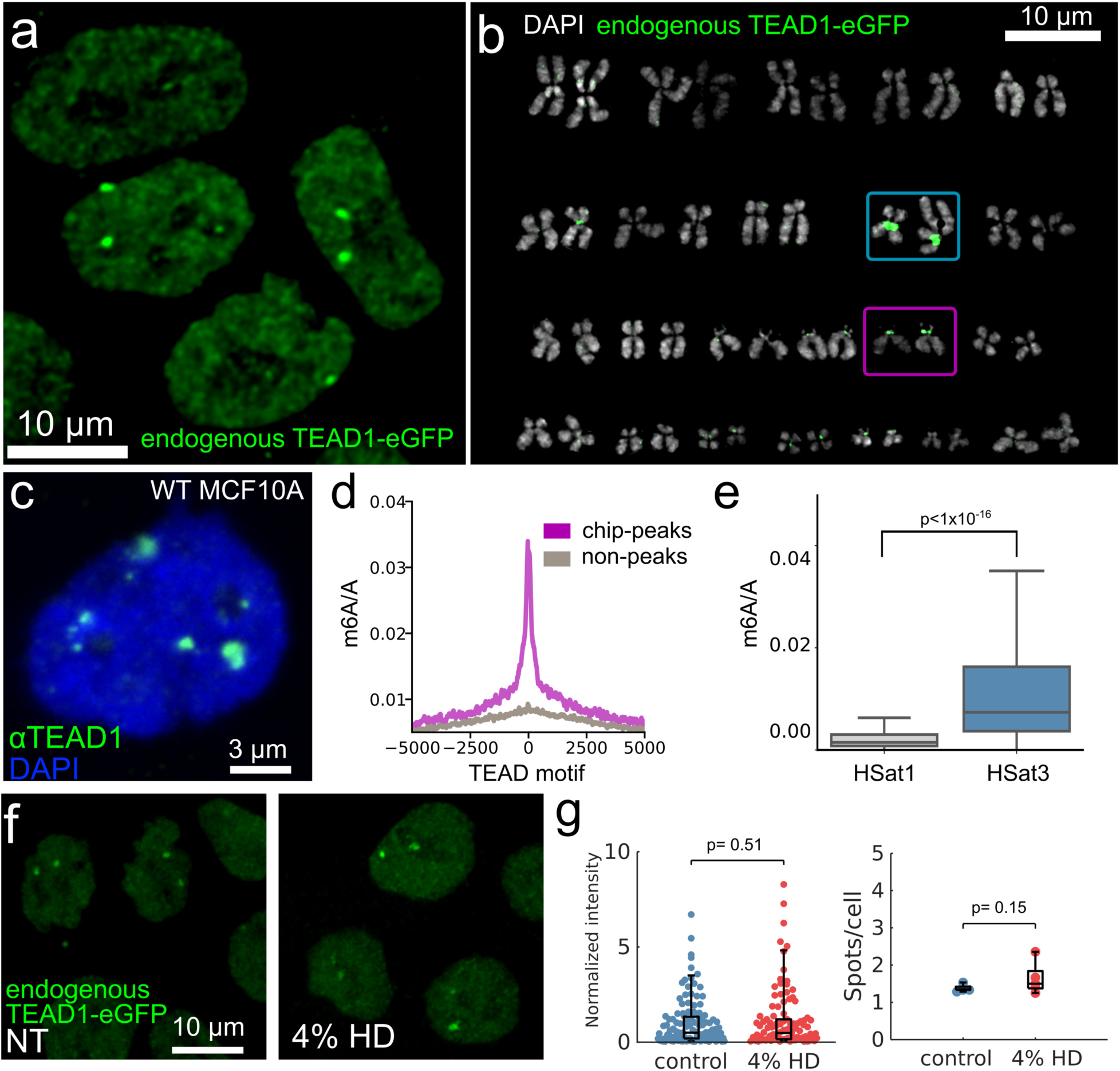
Endogenous TEAD1 binds HSat3. (a) TEAD1-eGFP knock-in (KI) at the endogenous locus in MCF10A cells shows clear sub-nuclear foci in live-cell microscopy. Image is a representative, single z-plane. (b) Representative metaphase spread of MCF10A with TEAD1-eGFP knock-in stained with GFP nanobody. The karyogram shows localization to the two largest HSat3 arrays (blue=chromosome 9, purple box=chromosome 15). To preserve TEAD1 signal, no other FISH or DNA stains could be used to generate banding patterns, thus chromosomes are organized by size as an approximation for the karyotype. MCF10A has a known translocation on chromosome 9^40^, showing two distinct chromosome sizes. Consistent results were obtained in two other metaphase spreads. (c) Immunofluorescence for TEAD1 in WT MCF10A, showing untagged endogenous TEAD1 shows sub-nuclear foci in unfixed wildtype (WT) MCF10A nuclei isolated for DiMeLo-seq. Image is a representative single z-plane. (d) TEAD1-targeted DiMeLo-seq in WT MCF10A. m6A signal is normalized to A, and centered at TEAD1 motifs within known TEAD1 ChIP-seq peaks. As an internal control, the signal is plotted at 25,000 randomly selected TEAD1 motifs outside of known ChIP-seq peaks and not within HSat3. (e) Distribution of m6A/A per read aligned to either HSat1 (N=668 reads) or HSat3 (N=1961 reads), showing specificity of TEAD1 binding to HSat3, as HSat1 is not enriched for TEAD motifs. The box plot lines are as follows: center line, median; box limits, upper and lower quartiles; whiskers, 1.5x interquartile range. A one-tailed Wilcoxon rank sum test was performed to compare m6A/A enrichment between HSat1 and HSat3. (f) Live-cell microscopy of endogenous TEAD1-eGFP in MCF10A in untreated and 10 minutes 4% 1,6-Hexanediol (HD) treatment showing no change in foci intensity or frequency. (g) Quantification of data from panel f. For the foci intensity comparison, Nfoci(control)=117, Nfoci(HD)=106. For number of foci per cell comparison: Ncells(control)=172, Ncells(HD) =141, with individual data points representing the mean for each region of interest. Image is a representative single z-plane. The experiment was performed twice, with similar results; data shown are from one biological replicate.

Both chromosome 9 copies showed pericentromeric staining, corresponding to HSat3 (MCF10A cells have a known chromosome 9 translocation, causing one copy to be longer^40^). Further, there is clear TEAD1 signal on many of the HSat3-containing acrocentric chromosomes (13-15, 21-22). Chromosome 15 in particular has an 8 Mb HSat3 array, corresponding to the brightest TEAD1 signal on the acrocentrics. Finally, we performed immunofluorescence staining on unfixed, permeabilized, wildtype MCF10A nuclei using an anti-TEAD1 primary antibody and imaged using confocal microscopy. These nuclei showed bright TEAD1 foci (Figure 4c), suggesting that the TEAD1-eGFP tag used in other experiments does not introduce localization artifacts, and that TEAD1 remains strongly bound to HSat3 without fixation.

We further validated endogenous TEAD1 binding to HSat3 using anti-TEAD1 DiMeLo-seq in unfixed, wildtype MCF10A cells, yielding m6A enrichment at known TEAD1 binding sites (Figure 4d). Within the sample, we measured bulk enrichment of m6A/A in HSat3 compared to an average of HSat1-A and -B, which are distinct satellite repeat families also present on acrocentric chromosomes and not predicted to be bound by TEAD (Figure 4e).

Phase separation of TFs into condensates has been reported for various TFs, including endogenous YAP and TEAD1^41^. Because phase separation is highly dependent on concentration, the high-frequency motifs embedded in satellite DNA may seed phase-separation through increased local concentration, as proposed for HSF1^42^. To test whether the observed TEAD binding to HSat3 is driven by phase separation, we incubated cells with 4% 1,6-Hexanediol, but this did not dissolve endogenous TEAD1-eGFP foci in live cells or reduce foci intensity (Figure 4f-g). In summary, our data suggest that endogenous TEAD1 binds HSat3 primarily through motif recognition.

### YAP localization to HSat3-bound TEAD depends on cellular state

TEAD-family factors require co-activators or co-repressors to regulate gene transcription. YAP is the best-characterized co-activator of TEADs. The YAP/TEAD complex mostly functions at gene enhancers, driving a transcription program that promotes cell proliferation^43^. Because TEAD1 appeared constitutively bound to HSat3, and YAP distributes well into the nucleus in interphase MCF10A^44^, we hypothesized that YAP should readily bind TEAD1/HSat3 foci. Using previously generated MCF10A YAP-eGFP/TEAD1-mCherry KI cells^44^, we did not observe increased YAP signal corresponding to the large TEAD1-HSat3 foci in interphase cells using 60X confocal microscopy (Figure 5a,c). However, mitotic cells showed strong YAP colocalization to TEAD1 foci (Figure 5b-c). Thus, the YAP/TEAD/HSat3 complex can form, but the formation of large YAP foci is cell-state dependent.

**Figure 5:**
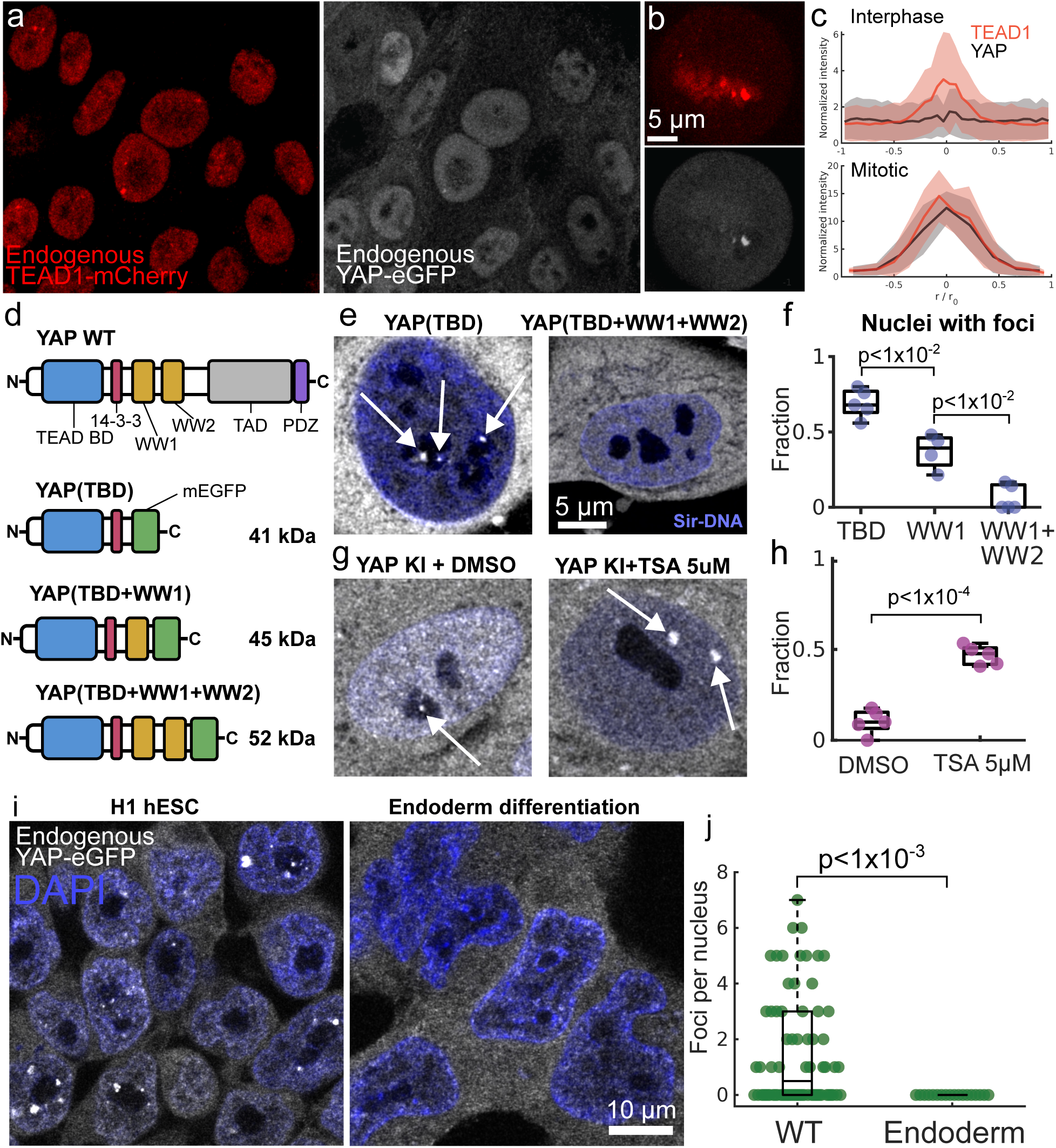
Cell-state dependent regulation of YAP binding to TEAD1/HSat3. (a-b) Two-color imaging of endogenous TEAD1-mCherry and YAP-eGFP shows no significant colocalization at HSat3 foci in interphase cells, and strong colocalization in mitotic cells. (c) Intensity analysis of images from panels a-b. For interphase cells, The TEAD1-mCherry intensity profile was generated from 23 foci detected from 55 cells. The YAP-eGFP intensity profile was generated by querying the TEAD1 foci pixels in the GFP channel. For mitosis, a representative cell was used to generate the intensity profile using the brightest TEAD1 focus. Shaded region represents the standard deviation. Experiment was performed three times with similar results. (d) Diagrams of YAP protein domains. Blue: TEAD binding domain (TBD), Red: 14-3­3 binding domain, Yellow: WW1 and WW2 domains, grey: transcription activation domain (TAD), purple: PDZ domain. YAP truncations fused to mEGFP and their molecular weights are shown. (e-f) live-cell super-resolution imaging shows frequent YAP(TBD) sub-nuclear foci which are nearly absent in YAP(TBD+WW1+WW2). Cells were stained with Sir-DNA for a live-cell nuclear marker. The fraction of cells with at least one detected focus are shown in (f). The number of cells per condition are as follows: N(TBD)=130, N(WW1)=90, N(WW1+WW2) = 52. Individual data points represent mean per ROI. Significance was assessed by one-tailed t-tests. Repeated two to three times with similar results. (g) Live-cell microscopy of endogenous (i.e. full length) YAP-eGFP knock-in (KI) treated with 5 μM Trichostatin A or DMSO, and stained with Sir-DNA. Treated cells show YAP localization to large sub-nuclear foci. (h) The fraction of cells with at least one detected sub-nuclear focus. The number of cells per condition are as follows: N(DMSO)=165, N(TSA)=106. Individual data points represent mean per ROI. Significance was assessed by a one-tailed t-test. Repeated twice with similar results. (i) Live-cell microscopy of H1 hESCs with endogenous YAP-eGFP stained with Sir-DNA. Untreated H1 cells frequently show many YAP foci which are immediately lost upon Activin A treatment to promote endoderm differentiation. (j) The foci count per cell nucleus is plotted for WT and endoderm differentiated cells. The number of cells per condition are as follows: N(WT) =72, N(endoderm)=18. A one-tailed Wilcoxon rank sum test was performed to assess statistical significance between conditions. Experiment performed twice with similar results; data shown is from one biological replicate.

To better understand regulation of YAP/TEAD on HSat3, we generated a series of YAP truncations fused to mEGFP containing the YAP TEAD-binding domain (TBD) (Figure 5d). Because endogenous YAP was not readily localized to TEAD1 in interphase cells, we employed live-cell super-resolution imaging with Zeiss Airyscan to assess YAP(TBD) binding to TEAD/HSat3 foci. We observed frequent localization of YAP-TBD-meGFP to sub-nuclear foci (Figure 5e-f). Whereas both YAP(TBD+WW1) or YAP(TBD+WW1+WW2) decreased the frequency of sub-nuclear foci, with almost no cells showing detectable YAP(TBD+WW1+WW2) foci (Figure 5e-f). These results suggest the domains of YAP may regulate its ability to bind TEAD1/HSat3, as the WW domains are responsible for multiple protein-protein interactions^45^.

Given the known heterochromatic status of HSat3, we hypothesized that treatment with Trichostatin A (TSA), a potent HDAC class I/II inhibitor, would de-condense HSat3 heterochromatin. Treatment of MCF10A cells with TSA led to a dramatic increase in YAP localization to sub-nuclear foci (Figure 5g-h). To our surprise, we found small nucleolar YAP foci in untreated cells using Airyscan (Figure 5g).

We thus hypothesized that endogenous YAP/TEAD1 binding to HSat3 does occur in interphase MCF10A, but foci were not detectable using 60X confocal microscopy. Although YAP is generally excluded from the nucleolus, we observed frequent nucleolar foci (Supplementary Figure 14). To prove these are indeed TEAD-bound HSat3, we generated a dual knock-in of endogenous TEAD1-HALO and YAP-eGFP in MCF10A. Indeed, nucleolar foci always colocalized with TEAD1-HALO stained with HALO-tag JF647 dye (Supplementary Figure 15), and all TEAD-family members colocalize 1:1 with ZFsat3, including foci within the nucleolus (Supplementary Figure 8). This is consistent with the genome organization of rDNA-containing acrocentric chromosomes, where HSat3 arrays are found on both sides of the nucleolar organizing regions (i.e. rDNA repeats plus distal- and proximal-junctions) (Supplementary Figure 16).

The presence of non-ribosomal DNA localization within the nucleolus has not been thoroughly explored. For example, we found only one decades-old report describing satellite DNA inside the nucleolus using DNA FISH^46^, whereas previous work typically describes satellite DNA as perinucleolar^47^. This lack of prior observations may be due to fixation artifacts and limits of resolution, as live-cell, super-resolution imaging of satellite DNA localization has not previously been reported, to the best of our knowledge.

Reports using dCas9 have shown the large chromosome 9 loci^25^, but not the smaller HSat3 arrays shown in our study.

Embryonic stem cell chromatin is largely euchromatic, whereas differentiation drives cell-type specific heterochromatin formation. This trajectory is also true for repetitive elements^48^. We next asked if HSat3 was more physically accessible to YAP/TEAD binding in hESCs. Using endogenous YAP-eGFP KI in H1 hESCs, we observed persistent large nuclear YAP foci in most interphase cells (Figure 5i-j). Whereas definitive endoderm (DE) differentiation led to immediate loss of detectable nuclear YAP foci (Figure 5i-j), despite ample nuclear YAP localization.

In summary, our results imply that the heterochromatic nature of HSat3 decreases YAP/TEAD interaction on HSat3. However, small nucleolar TEAD/HSat3 arrays are readily accessible to YAP in terminally differentiated MCF10A cells. Surprisingly, mitotic chromosomes permit ample YAP/TEAD/HSat3 interactions, despite being in a condensed state. This may be explained by multiple reports of chromatin accessibility in mitosis where TFs ‘bookmark’ gene promoters and enhancers, despite the large-scale condensation of chromatids^49,50^. Previous work showed fast and strong silencing of rRNA transcription following DE differentiation of hESCs^51^, which correlates well with the observed loss of nuclear YAP foci in hESCs upon DE differentiation. We thus hypothesized that HSat3 adjacent to rDNAs may regulate RNA Polymerase I (Pol-I) transcription via YAP/TEAD.

Exogenous expression of YAP and the paralogue TAZ has previously been reported to induce compartmentalization of TEAD-family factors, whereas TEAD did not localize to phase-separated compartments on its own^52,53^. However, a recent report demonstrated the localization of FLAG-TEAD4 to sub-nuclear compartments. These inconsistent observations may be due to immunofluorescence and/or fixation artifacts, as unfixed immunofluorescence showed consistent sub-nuclear endogenous TEAD1 foci (Figure 4c). Our results highlight the need to endogenously tag YAP and TEAD for live-cell imaging to avoid overexpression and immunofluorescence artifacts.

### YAP/TEAD/HSat3 regulate rDNA transcription in the nucleolus

The Hippo pathway acts as a global regulator of cell growth by sensing intracellular and extracellular cues, including GPCR-ligands, extracellular matrix mechanics, temperature, and metabolic states^16^. This pathway shows sensitivity to signaling inputs on a timescale of minutes^44,54,55^, providing a rheostat-like mechanism for regulating cell proliferation. Ribosome biogenesis represents a massive energy demand for the cell. rRNA genes are the most highly transcribed genes, representing 80-90% of total RNA in cells^56^.

In humans, not only are all rDNA arrays surrounded by multiple HSat3 arrays (Figure 1a), the homologues of human chromosomes 9 and Y in orangutans contain rDNA arrays adjacent to HSat3 (Supplementary Figure 16)^9,57^. Further, all rDNA-containing chromosomes in great apes contain nearby HSat3 arrays (Supplementary Figure 16), suggesting a well-conserved relationship between these two elements. Given the known roles of the Hippo pathway, and localization of YAP/TEAD to HSat3, we hypothesized that YAP/TEAD activity may regulate rRNA synthesis. Although the YAP/TEAD complex is only known as a RNA Polymerase II (Pol-II) regulator, many canonical Pol-II TFs directly regulate rDNA transcription, including cMYC^58^ and CEBPɑ^59^.

In MCF7 TEAD1-4 knock-out cells^60^, we found a 31% reduction in total RNA/DNA compared to wildtype MCF7 cells, suggesting that YAP/TEAD activity is upstream of rRNA transcription (Figure 6a). However, this effect could be due to global loss of YAP/TEAD activity, not necessarily through direct regulation of RNA Pol-I activity.

**Figure 6:**
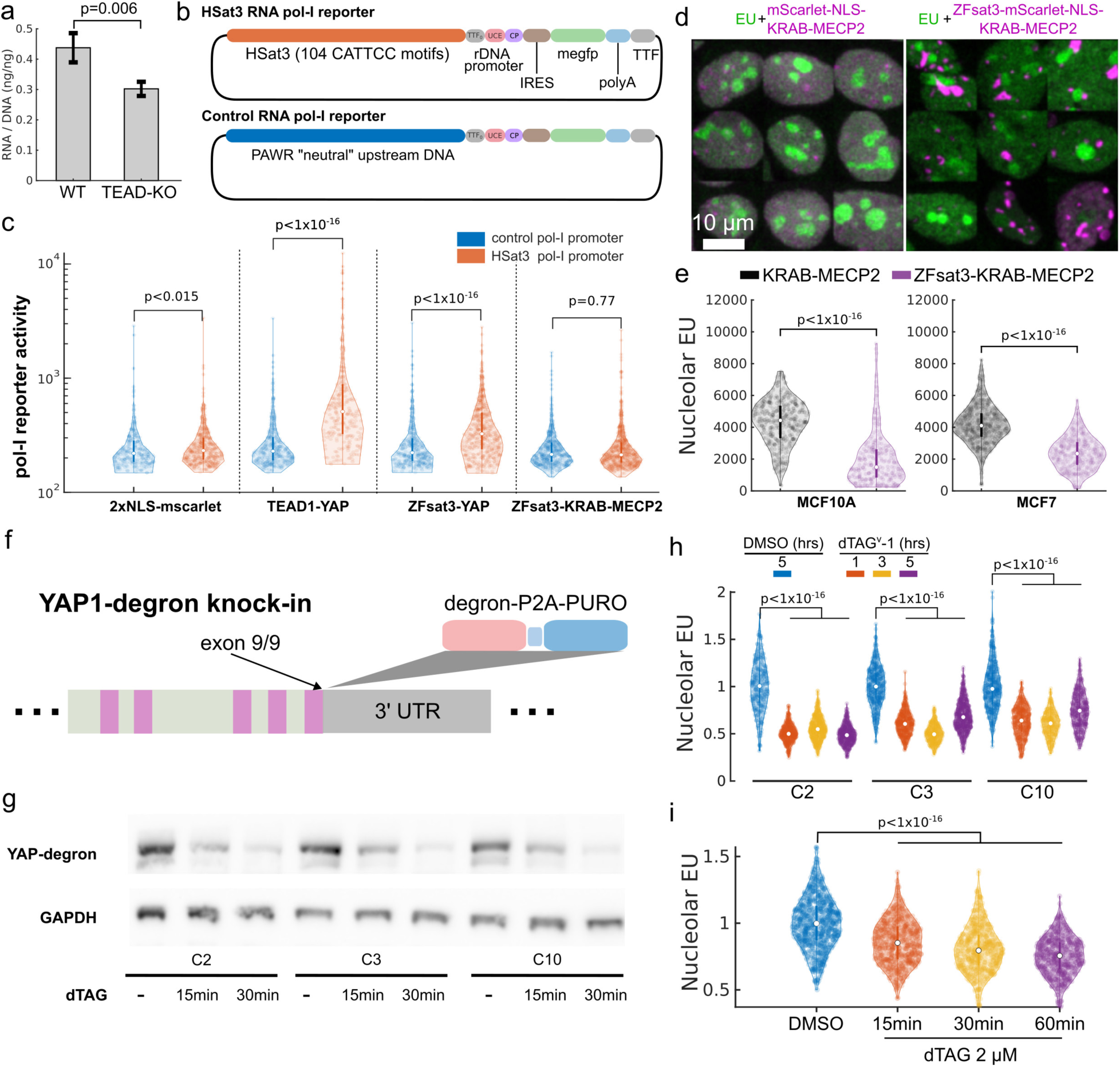
YAP positively regulates RNA Pol-I activity. (a) Permanent TEAD KO in MCF7 cells led to reduced RNA/DNA ratios. Bar plot generated from 3 biological replicates. Bar plot error represents standard deviation. Statistical significance was assessed using a one-tailed T-Test. (b) Design of RNA Pol-I reporter constructs with either HSat3 elements containing 104 CATTCC sites, or an equivalent-length DNA segment from the human PAWR gene. (c) Results of RNA Pol-I reporter assays. 293T cells were transfected with either control or HSat3-Pol-I reporters and 2xNLS-mScarlet (non-activating), TEAD1-mScarlet-YAP fusion, ZFsat3-mScarlet-YAP fusion, or ZFsat3-mScarlet-KRAB-MECP2. Cells were segmented using the nuclear mScarlet, and GFP signal was quantified from the cytoplasm of segmented cells. The number of cells per condition are as follows: N(control+NLS-mscarlet)=270, N(HSat3+NLS-mscarlet)=496, N(control+TEAD-YAP)=369, N(HSat3+TEAD-YAP)=275, N(control+ZFsat3-YAP)=358, N(HSat3+ZFsat3-YAP)=561, N(control+ZFsat3-KRAB-MECP2)=514, N(HSat3+ZFsat3-KRAB-MECP2)=720. A one-tailed Wilcoxon rank sum test was performed to assess statistical significance between control and HSat3 reporter activity. (d) representative confocal microscopy images of EU assay with MCF10A cells expressing either mScarlet-NLS-KRAB-MECP2 (KM) or ZFsat3-mScarlet-NLS-KM (ZFsat3-KM). Image is constructed by tiling 9 representative nuclei within each condition. (e) Quantification of nucleolar EU fluorescence intensity in MCF10A and MCF7 cells stably expressing ZFsat3-mScarlet-KM and mScarlet-KM. The number of nucleoli per condition are as follows: N(MCF10A KRAB-MECP2)=447, N(MCF10A ZFsat3-KRAB-MECP2)=270, N(MCF7 KRAB-MECP2)=1729, N(MCF7 ZFsat3-KRAB-MECP2)=945. A one-tailed Wilcoxon rank sum test was performed to assess statistical significance between KM and ZFsat3-KM. (f) Diagram depicting knock-in of degron-p2a-puromycin cassette at the human YAP1 locus. (h) 1 hr EU incorporation assay for H1 hESC YAP-degron clonal cell lines treated with DMSO or 2 μM dTAGv1 at various times. The number of nucleoli per condition are as follows: N(DMSO C2)=277, N(dTAG 1hr C2)=446, N(dTAG 3hr C2)=557, N(dTAG 5hr C2)=477, N(DMSO C3)=446, N(dTAG 1hr C3)=494, N(dTAG 3hr C3)=393, N(dTAG 5hr C3)=629, N(DMSO C10)=485, N(dTAG 1hr C10)=484, N(dTAG 3hr C10)=346, N(dTAG 5hr C10)=414. (i) 1hr EU incorporation assay for H1 hESC YAP-degron clone 2 after 15, 30, and 60 min dTAGv1 treatment. N(DMSO)=854, N(dTAG 15 min)=577, N(dTAG 30 min)=560, N(dTAG 60 min)=548. Data represents one biological replicate. The experiment was performed twice with similar results.

We hypothesized that HSat3 arrays act as enhancer-like elements for RNA Pol-I transcription via YAP/TEAD. To test this, we generated a RNA Pol-I enhancer construct consisting of the rDNA Upstream Control Element (UCE) and core promoter, followed by an IRES-meGFP-polyA expression cassette, and downstream Transcription terminator 1 (TTF1) sites (Figure 6b). To test the effects of HSat3 on RNA Pol-I transcription, we inserted either a 1.8 kb HSat3 element containing 104 GGAATG motifs (i.e. canonical TEAD binding sites), or a 1.8 kb neutral element from the human PAWR gene devoid of TEAD binding sites ahead of the rDNA promoter. The reporter was sensitive to both actinomycin D and an RNA Pol-I inhibitor Bmh-21 (Supplementary Figures 17-18). Co-transfecting the reporter and a 2xNLS-mScarlet expression plasmid showed a minor change in activity between the control and HSat3 rDNA reporters (Figure 6c). However, co-transfecting a chimera gene consisting of the TEAD1 DNA binding domain and the transcription activating domain (TAD) of YAP (TEAD-YAP) strongly increased the activity of the HSat3 rDNA reporter, but not the control rDNA promoter (Figure 6c). Similarly, co-transfecting ZFsat3-YAP activated the HSat3 rDNA reporter, but to a lesser extent. This is potentially due to reduced stoichiometry as each ZFsat3 targets 18 bp, compared to the TEAD1 footprint which is closer to 6 bp. To test the specificity of YAP activation of RNA Pol-I, we generated a TEAD1 fusion to VP64, a generic potent RNA Pol-II activator. TEAD1-VP64 did not activate the rDNA promoter, demonstrating the YAP transcription activation domain is specifically activating the RNA Pol-I reporter (Supplementary Figure 19).

The genetic deletion of all rDNA-adjacent HSat3 arrays cannot be done due to the sheer number of individual arrays (Supplementary Figure 1). Further, compensatory mechanisms may develop during clonal selection of HSat3-deleted cells. Therefore, we targeted HSat3 directly with a KRAB-MECP2 (KM) repressor^61^ fused to ZFsat3-mscarlet (ZFsat3-KM). The transcription rate of rRNA can be accurately quantified using incorporation of an ethynyl modified uridine analog (5-Ethynyluridine, EU), which shows strong nucleolar signal proportional to rRNA transcription^62^. In both MCF10A and MCF7, ZFsat3-KM showed strong inhibition of rRNA transcription compared to expression of KM alone (Figure 6d-e) or ZFsat3-mScarlet alone (Supplementary Figure 20). In some cases with high ZFsat3-KRAB-MECP2 expression, cells showed almost no incorporation of EU in nucleoli (Figure 6d). This result suggests that the chromatin status of HSat3 is important for normal rRNA transcription.

We surmised the connection between rRNA transcription and YAP/TEAD/HSat3 is strongest in hESCs, given the ample localization of YAP to nuclear foci in most cells (Figure 5i). We generated clonal hESC lines with homozygous endogenous YAP-degron KI (Figure 6f, Supplementary Figure 21), allowing acute degradation of YAP (Figure 6g). All clones showed strong reduction of nucleolar EU signal following 1, 3, and 5 hr degradation of YAP followed by 1 hr EU incorporation (Figure 6h). We further probed EU incorporation after 15, 30, and 60 minutes of YAP degradation for clone 2. This showed detectable EU reduction even after 15 minutes of YAP degradation followed by 1 hr of EU incorporation (Figure 6i). We measured a ∼25% drop in rRNA transcription rate after 1 hr of YAP degradation plus 1 hr EU incorporation, however, it is possible that the YAP paralogue TAZ or other HSat3-bound TFs are compensating for loss of YAP. In comparison, degradation of degron-tagged CEBPα, which directly binds the rDNA promoter, showed a ∼7% decrease in rDNA synthesis after 2 hours of dTAGv1 treatment, measured using FISH against nascent rRNA^59^. The speed and magnitude of reduction in rDNA synthesis after acute degradation of YAP implies a direct relationship, although second-order effects cannot be formally ruled out.

Because HSat3 arrays are found adjacent to all rDNA arrays, we hypothesized that YAP/TEAD/HSat3 may be co-localized with UBTF, a core rDNA TF that marks active rDNAs (Figure 7a). Two-color imaging of mScarlet-UBTF and endogenous YAP-eGFP in H1 hESCs showed 58% of YAP foci contacting UBTF foci within nucleoli (Figure 7b).

**Figure 7:**
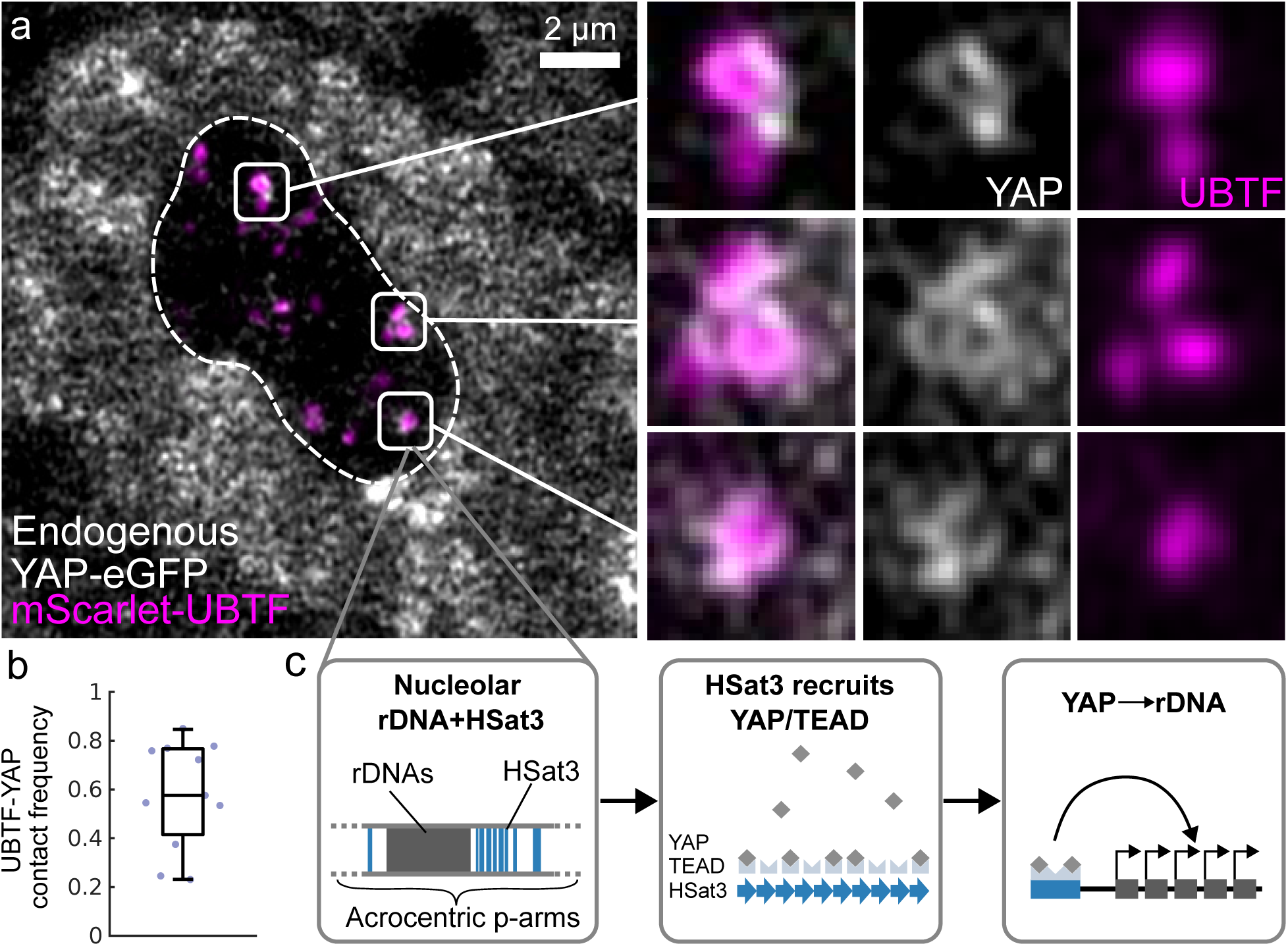
Colocalization of YAP and UBTF in nucleoli. (a) Live-cell super-resolution imaging (Airyscan) of mScarlet-UBTF and endogenous YAP-eGFP in H1 hESCs. Dashed line represents the outline of the nucleolus. Insets show contact between UBTF foci and YAP foci. (b) Nucleolar UBTF and YAP foci are segmented, and the fraction of UBTF foci that touch YAP foci is quantified per cell. Data points represent mean contact frequence per cell (N=11 cells, with a total of (c) Summary of the major findings presented in this work. Left: Detection of nucleolar HSat3. Center: endogenous TEAD binds HSat3 and recruits YAP. Right: YAP regulates RNA Pol-I activity.

Together, these results show YAP/TEAD/HSat3 not only co-localizes with rDNA machinery in live cells, but it can directly activate the rDNA promoter (Figure 7c). Thus, we propose two possible models: (1) The YAP/TEAD complex on HSat3 forms 3D interactions with rDNA in the nucleolus, or (2) TEAD/HSat3 increases the nucleolar concentration of YAP, allowing more frequent contacts with the rDNA promoter.

## DISCUSSION

Our work has unveiled a set of novel HSat3-binding TFs, many of which have potent transcription driving functions. These TFs function in various signaling contexts, including development, immune response, T-cell activation, and human cancers. Our findings suggest that constitutive heterochromatin contains a diverse and dynamic collection of proteins^63,64^, rather than being an inert or purely repressive monolith^65^. Characterizing the complete set of satellite binding TFs and their functions may add to this growing body of previously unappreciated protein-DNA interactions.

We further explored the potential biological functions of our findings by focusing on TEAD, which is widely expressed as part of the Hippo pathway and normally plays a role in cell proliferation. We observed HSat3-bound TEAD foci within nucleoli, along with TEAD’s co-activator YAP co-localized with the RNA Pol-I transcription factor UBTF. We thus hypothesized that YAP/TEAD may play a direct role in rDNA regulation, mediated by binding to HSat3, given the well-documented functions of the Hippo pathway as a growth control network. Acute degradation of YAP revealed a rapid decline in endogenous rDNA transcription, and the YAP transcription activation domain directly activated an RNA pol-I reporter, confirming that YAP can promote RNA Pol-I activity in cis. Lastly, recruitment of a KRAB-MECP2 repressor to HSat3 led to strong inhibition of rDNA transcription, confirming that factors bound to HSat3 can influence RNA Pol-I activity.

Because HSat3 arrays and rDNA arrays are often hundreds of kilobases apart, HSat3-bound YAP/TEAD and rDNAs may directly interact via physical proximity within the nucleolus. Alternatively, because co-activator binding is transient and dynamic, HSat3-bound TEAD may simply increase the local concentration of the co-activator YAP, enhancing rDNA expression at a distance. Future experiments will need to resolve this question and further explore the precise mechanism by which YAP modulates rDNA activity. Improved genetic tools may allow for the mutation/deletion of the satellite DNA itself, though this remains extremely challenging given the large number and size of satellite DNA arrays across many chromosomes throughout the genome.

To the best of our knowledge, human satellite DNAs, TEAD, and YAP have not previously been observed inside the nucleolus, likely due to a lack of suitable super-resolution live-cell imaging assays. A recent report described sub-nuclear TEAD4 condensates in live-cell imaging, which are dramatically enriched following glucose starvation^66^. Although the genomic basis of these foci was not investigated, our results suggest these are indeed HSat3-bound TEAD. We predict that the YAP paralogue TAZ may operate on HSat3-bound TEAD in a similar manner to YAP. Because TAZ has overlapping and distinct expression and cellular functions ^67,68^, further investigations in cell types with strong TAZ activity may provide insights into the differential regulation of rDNA by YAP and TAZ. Hypothetically, the relationship between satellite DNA and rDNA regulation may extend beyond HSat3 and YAP/TEAD. Here, we propose a model in which upstream regulators of rDNA transcription are brought into the nucleolus via adjacent satellite repeats. Multiple IRF family members were previously found to directly bind the rDNA promoter in human and mouse cells^59^. In our work, we demonstrated IRF3 binding to HSat3, further supporting the notion that pools of rDNA-regulating TFs are locally enriched via satellite DNA repeats.

Large HSat3 arrays appear to have originated adjacent to rDNA, as they are found consistently on rDNA-containing chromosomes across great apes^9^. Human chromosomes 9 and Y have the largest amount of HSat3 per chromosome, yet no rDNA arrays. However, the homologous chromosomes in orangutan, the most distantly related great ape, do harbor rDNA arrays^9,57^. This raises interesting questions about the potential role of the HSat3-Hippo-rDNA regulatory axis in the evolution of great apes.

The recruitment of TFs like TEAD to HSat3 may change the underlying chromatin state. Future studies will need to explore the effects of these TFs on phenomena like chromatin compaction, histone modifications, HP1 association, and satellite transcription. If TF-loaded satellite arrays have a less repressive chromatin state, they may help to insulate highly active rDNAs from the surrounding repressive heterochromatin. Our work suggests that upstream activation of the Hippo pathway would reduce YAP-TEAD/HSat3 loading by YAP phosphorylation^16^, shifting HSat3 towards a more repressive state and reducing rDNA transcription.

YAP/TAZ are implicated in a variety of human cancers^16^. The high metabolic demand of tumor growth requires ample ribosome biogenesis, and thus rRNA transcription^69^. Further studies on oncogenic YAP/TAZ activation may reveal an underlying connection to rRNA transcription via our proposed model. Overexpression of activated YAP induces transcription of ribosomal protein components^70^, and increases global translation even during serum starvation^71^. In line with our results, we propose that YAP promotes rRNA synthesis directly to meet increased translation demands induced by its own downstream activities.

In summary, we have identified multiple, highly conserved transcription factors that directly bind massive, poorly understood satellite repeats in the human genome. Our results suggest one potential function of these satellite arrays is to act as enormous gene regulatory elements that can modulate rDNA expression via TF enrichment in the nucleolus.

## METHODS

### Cell culture

293T: 293T cells (ATCC CRL-3216) were cultured in DMEM base media with 10% FBS, 1xPEN/STREP supplement. For transfections, Lipofectamine 3000 (Thermo Fisher Cat L3000001) was used according to the manufacturer’s instructions.

MCF10A: MCF10A cells were cultured as previously described^44^.

H1 hESC: H1 cells were obtained directly from WiCell at passage 22. Cells were cultured according to WiCell’s protocol using Matrigel coating (Corning 354277), mTESR Plus (Stem Cell Technologies 100-0276) and Versene EDTA (Fisher Scientific A4239101). Briefly, cell culture surfaces were coated with Matrigel for 2 hrs or longer at 37 C in the tissue culture incubator. For passaging, cells were rinsed with once Versene, incubated with Versene for 8.5 minutes at room temperature. Versene was aspirated, and then fresh mTESR Plus media was used to lift hESC colonies from the dish. Endoderm differentiation was carried out by washing once with DPBS, and culturing with 50 ng/mL Activin A (Gibco Cat. No. PHC9564) with 3μM CHIR99021 (Selleck Chemicals Cat. No. S2924) in RPMI 1640 base media, replacing media once after 24 hrs.

Stem cell Ethics statement: H1 hESC’s were cultured in vitro in accordance with the Stem Cell Research Oversight Panel (SCRO) which complies with federal, state and Stanford policies.

For transient transfections, cells were passaged using Accutase (Stem Cell Technologies 07920) to generate single-cell suspensions, followed by seeding with Revitacell Supplement (Fisher Scientific A2644501). The next day, cells were transfected with Lipofectamine Stem Cell reagent (Invitrogen STEM00015) following manufacturers instructions.

### HSat3 motif analysis

Karyoplot of HSat3 arrays in the human genome was made using the centromere/satellite annotations of T2T-CHM13v2.0 and karyoplotteR package in R.

To calculate the enrichment of TF motifs across Hsat3, we used the MEME-Suite tool SEA: Simple Enrichment Analysis. Background sequences were generated from T2T-CHM13v2.0 reference genome using the 2000 bases upstream of 5000 randomly selected genes from NCBI ref seq annotation. For each selected gene, ten 50-base pair sub-strings were randomly chosen, providing 50,000 “reads” for background data. The input sequences were similarly generated from HSat3 annotations designated by sub-family^1^. Because HSat3 is inherently repetitive, we generated substrings with a mean length of 50 base pairs with standard deviation of 4 to avoid biasing. For each HSat3 element, the complete set of 50+/-4 base pair strings were used for SEA analysis. For TF motifs, we used the JASPAR 2024 database (https://jaspar.elixir.no/download/data/2024/CORE/JASPAR2024_CORE_non-redundant_pfms_meme.txt). A heat-map was generated using a custom python script, plotting the enrichment values from SEA. We clustered individual HSat3 arrays by subfamily (A1,B2,B6, etc) using Ward hierarchical clustering with Euclidean distance (scipy.cluster.hierarchy). Raw data and scripts will be available upon publication.

### ChIP-seq screen

We downloaded publicly available ChIP-seq data from the ENCODE portal to mine for putative HSat3-binders. The ENCODE cell lines included in this screen are: HepG2, WTC11, H1, HCT116, A549, GM23338, K562, HEK293, GM12878, MCF-7, HEK293T, SK-N-SH, and IMR-90. We screened a total of 2,407 ENCODE accessions, ensuring each had at least one experimental and one control dataset, with most accessions containing at least two replicates. Data were specifically split between male and female cell lines to account for sex-specific differences, and only accessions with paired control datasets were downloaded. This resulted in 13,542 FASTA files that were downloaded and aligned to the Chm13v2.0 genome with or without chromosome Y depending on cell line sex. Alignment was carried out using BWA^72^. We added an additional dataset for HSF1 ChIP-seq in Heat-shocked MCF7 cells as a positive control (GSE209667).

To calculate the enrichment ratio of transcription factor binding in satellite regions, we defined control regions as 2,000 randomly sampled windows of 100 kilobases each across the genome. For female cell lines, chromosome Y was excluded. The mean number of reads in these control regions was calculated for each sample and used as the denominator to normalize for antibody and ChIP biases inherent in the datasets.For each sample, both control and experimental, we calculated the ratio of reads in the satellite region to the mean reads in the control regions. To measure the effect size, we took the median of these ratios for all experimental datasets corresponding to a given cell line, transcription factor, and satellite DNA. This median was then divided by the median ratio obtained from the control datasets for the same sample parameters. Statistical significance was assessed using the Mann-Whitney U test to compare the effect sizes between control and experimental groups with the null hypothesis that the satellite regions have less or equal transcription factor binding compared to the control regions.

### Library of HSat3-binding TFs

Starting with a lentivirus vectors pHIV-EF1alpha-mcs-pgk-puro/hygro, we generated base constructs for N-spytag3-mScarlet-linker-X or X-linker-mScarlet-spytag3-C tagging of genes, where X is a restriction site for inserting arbitrary cDNAs via NEB HiFi assembly. Most TFs were sourced from the Multiplexed Overexpression of Regulatory Factors (MORF) Library, a gift from Feng Zhang (Addgene #1000000218). TFs not available within the library were amplified directly from cDNA. Base constructs, TF-expressing constructs, sequences and primers used to PCR TFs are found in Supplementary data (provided after publication).

TF cDNAs were PCR amplified using NEB Q5 2x mastermix (NEB Cat. No. M0541L) and gel-purified using Qiagen gel purification kit (Qiagen Cat. No. 28704). Assembly of each factor was done using NEB HiFi 2X mastermix (NEB Cat. No. E2621L) with 50 ng of digested backbone and 10-40ng of gel purified cDNA following standard protocol from NEB. Assembled constructs were transformed into NEB DH5alpha (NEB Cat. No. C2987H). All clones were validated by long read sequencing (Plasmidsaurus).

Lentivirus particles were generated for TF expression constructs using Pspax2 / pmd2.g (gift from Addgene). using 293T cells. Briefly, [transfection mix] in 6-well plates coated with poly-L-lysine (Millipore Cat. No. A005C). Virus containing supernatant was concentrated 100-fold using Lenti-X Concentrator (Takara bio Cat. No. 631231), and used immediately or frozen at −80 C.

293T cells were transduced with lentivirus and polybrene at 10 μg/mL (Millipore Cat. No. TR-1003-G), and selected using puromycin at 2.0ug/mL, or hygromycin B at 200 ug/mL.

### Design and validation of HSat3 Zinc Finger array

The original design of the HSat3 Zinc-Finger (ZFsat3) was done using a now defunct webtool (https://www.scripps.edu/barbas/zfdesign/designzf.php). However, the design principles are described in Wright et al^73^. An optimal protein sequence was obtained by targeting an 18-bp perfect HSat3 repeat target [5’-GGAAT-GGAAT-GGAAT-GGA-3’]. The resulting amino acid sequence including ZFs and linkers will be released upon publication.

The human codon optimized gene was generated by Twist Biosciences, and cloned into a pHIV backbone with an EF1 alpha promoter using restriction cloning. The ZFsat3 was cloned with a linker, mScarlet, and NLS at the C-terminus. Plasmid sequence available in the supplementary materials.

We validated the specificity of ZFsat3 binding to HSat3 using immuno-FISH. MCF10A cells were infected with lentivirus encoding ZFsat3-mScarlet-NLS, and selected using puromycin at 1μg/mL. After antibiotic selection, 20,000 cells were plated on Mattek imaging dishes (Mattek Cat. No. P35G-1.5-10-C). Dishes were pre-coated with human Fibronectin (Corning Cat. No. 356008) at 10 μg/mL for 1 hr at room temperature. Cells were rinsed once in DPBS, and fixed with 4% paraformaldehyde in DPBS for 15 minutes at room temperature. Fixed cells were permeabilized by three washes of 3 minutes each with 0.1% Triton X-100, and blocked for 1 hour with 3% BSA in PBS. Direct immunofluorescence was performed with anti-RFP antibody (Rockland Immunochemicals, Cat. No. 600-401-379) labeled with CF488 fluorescent dye using Biotium Mix-N-Stain (Biotium Cat. No. #92233), at 0.57 μg/mL for 1 hour at room temperature. Samples were then washed three times for 5 minutes in PBS, followed by post-IF fixation with 1% PFA for 10 minutes and three 1 minute PBS washes, and 10 minute aldehyde quenching with 20mM glycine in PBS.

For HSat3 DNA FISH, the samples were then incubated in 25% glycerol in PBS for 1 hour at room temperature, followed by three liquid nitrogen snap freeze-thaw cycles. Typical FISH protocols require 0.1 M HCl treatment to improve FISH probe penetration, which would bleach the dye-labeled RFP antibody. Instead, samples were lightly digested with DNAse-I (Ambion Cat. No. AM2222) at 0.5 U/mL for 5 minutes at room temperature followed by inactivation with EDTA at 2mM final concentration. The buffer was then exchanged to 50% formamide with 2X SSC (Invitrogen 20X SSC Cat. No. AM9763) for 1 hour at room temperature. Hybridization buffer (Empire Genomics Cat. No. 16250) with or without HSat3 probe at 0.4 ng / μL was added to the samples, followed by heat denaturation in an oven and overnight hybridization: 5 minutes 83C, 5 minutes 37C, 10 minutes 83C, and 37 C overnight in a humid chamber. Samples were then washed three times for 5 minutes in 2x SSC at 37 C, and three times in 0.1x SSC at room temperature. Finally, nuclei were stained with DAPI at 1 μg / mL for 10 minutes at room temperature, followed by a PBS rinse. Control samples showed loss of mScarlet fluorescence after FISH treatment, and strong signal for HSat3 without immunofluorescence staining of ZFsat3-mScarlet.

The HSat3 probe sequence is: “ATT+ CCA+ TTC+ CAT+ TCC+ ATT+ CC”, where “+” bases are linked nucleic acids. The probe was labeled with Alexa 594 at the 5’ end, and purified by HPLC (Integrated DNA Technologies, custom order).

### HSat3 colocalization assay

The imaging screen of HSat3 binding factors was carried out by transient transfection in 293T cells in 96-well imaging plates with 400uL of media per cell (Ibidi Cat.No:89626). The day before transfection 30k cells were plated per well. On the day of transfection, each well was transfected using Lipofectamine 3000 (Thermo Fisher Cat L3000001) following the manufacturer’s protocol. For single color imaging, 200 ng of the HSat3 TF was transfected. For two-color colocalization, 60ng ZFsat3-megfp and 180 ng of the HSat3 TF was transfected. 8 hrs post transfection, cell media was washed once.

### DiMeLo-seq

Samples were prepared following the v2 protocol (dx.doi.org/10.17504/protocols.io.b2u8qezw) with the following modifications: Reagents for the following experiment were freshly prepared, syringe filtered through a 0.2-µm filter, and then placed on ice. Cells were grown in 75 cm2 flasks at 37°C in 5% CO2, with 4 million live cells used per sample. Cells were pelleted at 500g for 3 minutes at 4°C and washed with PBS. To isolate nuclei, cells were resuspended in 1 mL of Dig-Wash Buffer (0.02% digitonin, 20 mM HEPES-potassium hydroxide buffer, pH 7.5, 150 mM sodium chloride, 0.5 mM spermidine, 1 Roche cOmplete EDTA-free tablet (11873580001) per 50 ml buffer and 0.1% BSA) and incubated on ice for 5 minutes and subsequently spun down at 500g for 3 minutes at 4°C. All consecutive spins were performed with the same conditions using wide bore tips for pipetting. The nuclei pellet was resolved in 75 µl of Tween-Wash Buffer (0.1% Tween-20, 20 mM HEPES-potassium hydroxide, pH 7.5, 150 mM sodium chloride, 0.5 mM spermidine, 1 Roche cOmplete EDTA-free tablet per 50 ml buffer and 0.1% BSA) containing primary antibody at a 1:50 dilution and then placed on a rotator at 4°C overnight. For exogenous expression of TEAD1-mscarlet in 293T cells, we used anti-RFP primary antibody (Rockland Immunochemicals, Cat. No. 600-401-379). For endogenous TEAD1 in MCF10A, we used anti-TEAD1 rabbit (CST Cat. No. 12292S).

Genomic DNA was extracted using NEB Monarch gDNA Extraction Kit (NEB Cat no T3010S). 1.5 µg of purified DNA from each sample was input into library preparation using the Native Barcoding Kit 24 V14 (Oxford Nanopore Technologies cat. no. SQK-NBD114.24). The library preparation was carried out according to the manufacturer’s protocol, with the following modifications: End-repair incubation time was increased to 10 minutes. A total of 221.5 ng of end-repaired DNA was loaded into the barcode ligation. LFB was used for the final adapter ligation clean up. Then, approximately 38.4 fmoles of the final pooled library was loaded onto the sequencer.

Sequencing was performed on a PromethION 2 Solo for 2-3 days using R10.4.1 flow cells.

All Pod5 files generated during sequencing were merged into a single Pod5 file using the pod5 merge command from the Pod5 package. Following this, Read IDs were extracted from the merged file using the pod5 view command. Once the Read IDs were obtained, alignment to the CHM13v2.0 genome was conducted. This process included DNA base calling, base modification calling, and demultiplexing utilizing dorado version 0.7.2 Linux x64. The resulting alignments were further merged, filtered, and indexed using samtools.

### DiMeLo-seq analysis

For analyzing m6A deposition centered at known binding sites, fibertools center was used to extract m6A base modifications centered at known ChIP-seq peaks defined by a bed file. Then reference matched base modifications were summed over a defined window centered at either ChIP-peaks or defined TF binding motifs within chip-peaks.

Known TF binding sites were collected from existing ChIP-seq data (listed below). Published narrowpeak files were first lifted over to the T2T-CHM13v2.0 (HS1) assembly using UCSC hgLiftOver (https://genome.ucsc.edu/cgi-bin/hgLiftOver). The resulting bed file was used to center ONT reads and quantify m6A modifications.

Chip-peak sources:

TEAD4: GSE66081

FOXA1: ENCODE (narrowPeak: ENCFF942VEL)

For quantifying deposition of m6A in satellite DNA, reads aligning to specified satellite regions were extracted using samtools view, then fibertools extract was used to identify modified bases on each read. We used a read length minimum of 2,000 bases, minimum mean base quality of 20 per read, and m6A minimum score 225 out 255.

### Metaphase spreads

Immunofluorescence of TEAD1-eGFP on metaphase nuclei using GFP-booster 488 (Chromotek Cat. No. gb2AF488) was performed following the protocol in Terrenoire et al^74^ with modifications. MCF10A TEAD1-eGFP knock-in cells were treated with 0.1 ug/mL colcemid overnight (∼15 hrs) to enrich mitotic nuclei. Cells were collected by trypsinization, resuspending in media, and spinning cells at 300g for 5 minutes. The cell pellet was resuspended in ice cold 75mM KCl at 4E5 cells/mL for 10 minutes on ice. After gently resuspending, cells were spun on a Shandon Cytospin 2 centrifuge, loaded at 200uL per funnel, at 1800 rpm for 10 minutes. Slides were then incubated in KCM buffer (120 mM KCl, 20 mM NaCl, 10 mM Tris/HCl pH 8.0, 0.5 mM EDTA, 0.1% Triton X-100) for 10 minutes at room temperature. Slides were incubated with GFP-booster 488 diluted 1:1000 in 3% BSA for 2 hours at room temperature, rinsed 3 times in PBS using, and mounted using Vectashield with DAPI (Vector Laboratories Cat. No. H-1200-10). Slides were imaged using 100X objective on a Delta Vision fluorescence microscope, followed by 3D deconvolution using default settings.

### Imaging of YAP-truncations

A Zeiss 980 Airyscan 2 was used to perform live-cell super-resolution imaging of various YAP-truncations stably expressed MCF10A cells. Lentiviruses were packaged as described above, and were used to tranduce WT MCF10A cells. Stable expression was achieved using puromycin at 1ug/mL. 40,000 cells were seeded on Mattek #1.5 imaging dishes (Mattek Cat. No. P35G-1.5-10-C) overnight. Prior to imaging, cell nuclei were stained with Sir-DNA for 1 hr at 1 μM, followed by a 5 min wash in warm media.

Samples were imaged live in an incubation unit at 37 C and 5% CO2. Two color imaging was performed, followed by 3D Airyscan image processing using default settings in Zeiss Zen software. Image segmentation is described below in “Microscopy segmentation and quantification”.

### Transcription reporter constructs

The RNA Pol-I reporter was inspired by Ghoshal et al^75^. The promoter, including the upstream binding enhancer and TTF1 binding sites through to the transcription start site was cloned into a PUC19 backbone, followed by an IRES, mEGFP, polyA, and downstream TTF1 sites to drive termination of Pol-I. Separately, a 2X copy of the pW1 HSat3 amplicon was cloned using restriction sites. The 2xpW1 (180Xbp) or an 1800bp region from the PAWR human gene was then inserted directly ahead of the Pol-I promoter.

The reporters were co-transfected with a control 2xNLS-mScarlet construct, or transgene known to bind HSat3. All co-transfected constructs were cloned into pHIV backbone driven by an EF1 promoter. To simply co-expression of a single activator, a TEAD1-YAP chimera was generated as follows from N to C-terminus: DNA-binding domain of TEAD1, linker, mScarlet, NLS, linker, and the C-terminus of YAP containing the transcription activation domain (TAD) or VP64 subcloned from Addgene plasmid # 61422. The 2xNLS control construct was generated by NEB HiFi assembly, with primers that encode the NLS’s. The ZFsat3-YAP(TAD) construct was generated by truncating full length YAP to just the WW+TAD domains.

### Transcription reporter assays

The activity from the RNA Pol-I reporters were assayed using transient transfection of 293T cells using Lipofectamine 3000 (Thermo Fisher Cat L3000001). 80,000 293T cells were seeded on 24-well Ibidi imaging plates (Ibidi Cat. No. 82426) overnight. The following day, cells were transfected with 300 ng of the reporter construct and 200 ng TEAD-YAP, ZFsat3-YAP, TEAD-VP64, ZFsat3-KRAB-MECP2, or 60 ng of 2xNLS-mScarlet in optimem.

After 6 hours, transfection reagent was removed, and replaced with cell culture media. Cells were imaged 24 hours after the start of transfection.

The specificity of this construct was tested using the RNA Pol-I inhibitors actinomycin-D at 50ng/μL and BMH-21 at 2 μM. Inhibitors were added after the initial 6 hour transfection period.

### CRISPR knock-in cell lines

CRISPR knock-in cell lines were generated using previously described methods^44^. In this work, the YAP-degron homologous recombination plasmid was generated by cloning the FKBP12 mutant domain into the YAP-eGFP-p2a-puro plasmid. For MCF10A, we used Viafect (Promega Cat. No. E4981), with a 8 μL reagent to 2 μg plasmid DNA (1.33 μg HR backbone, 0.67 μg px459 vector with YAP-KI sgRNA). H1 hESCs, we used Lipofectamine Stem Reagent (ThermoFisher Cat. No. STEM00003) with 5 μL reagent to 2.5 μg plasmid DNA (1.67 μg HR backbone, 0.83 μg px459 vector with YAP-KI sgRNA).

Cells were diluted and plated into a 96-well plate under puromycin selection for generating clonal cell lines. For H1 hESCs, we used 0.3 μg/mL puromycin selection, as cells were very sensitive to puromycin. We selected colonies with normal ESC morphology to assess the YAP-degron genotype using genomic PCRs and western blot (Supplementary Figure 21).

### Western blots

To confirm YAP-degron knock-in and rapid degradation, H1 YAP-degron clonal cells were seeded in 12-well tissue culture plates overnight following a 1:5 split ratio. Cultures were treated with DMSO, or 2 μM dTAG^v^-1 for 15 and 30 minutes in the tissue culture incubator. Samples were lysed in the plate using Laemmli SDS buffer (ThermoFisher Cat. No. J61337.AD) supplemented with 5% ꞵ-mercaptoethanol, and boiled for 5 minutes in a heat-block. Samples were run on a 4-12% Bis-Tris gel (Invitrogen Cat. No. NW04125BOX), with wet-transfer performed on a PVDF membrane. A 1 hour room temperature blocking was done with 2% Amersham ECL Prime Blocking Reagent (Cytiva Cat. No. RPN418) in Tris-buffered saline with 0.1% Tween 20. Membrane was cut to probe GAPDH (CST 2118S, 1:1000 dilution) and YAP (Santa Cruz sc-101199, 1:1000 dilution) in 2% ECL Prime in TBST overnight, followed by 3×5 minute washes in TBST. Rabbit/mouse conjugated secondary antibodies were incubated on membranes in 2% ECL Prime in TBST for 1 hour, followed by 3×5 minute washes in TBST. The membrane was imaged using a Gel-doc? Imager following incubation with Westernbright Quantum HRP substrate (Advansta Cat. No. K-12042-C20).

### EU assays

To drive targeted repression of HSat3, we fused ZFsat3-mScarlet to KRAB-MECP2^61^ sourced from dCas9-KRAB-MeCP2, a gift from Alejandro Chavez & George Church (Addgene plasmid # 110821; http://n2t.net/addgene:110821; RRID:Addgene_110821). As a negative control, we cloned a non-targeting construct mScarlet-KRAB-MECP2. Plasmids were assembled using restriction cloning (PCR primers in supplementary data, provided after publication).

Cells were seeded on 96-well imaging plates (Ibidi Cat.No:89626) at various densities to achieve 30-50% confluency on day of EU incorporation assay. For stable lentivirus transduced MCF10A lines (ZFsat3-mScarlet, ZFsat3-mScarlet-KRAB-MECP2, mScarlet), cells were seeded at 30k cells / cm^2^, and assayed 48 hrs post seeding. For YAP-degron time series, H1 hESCs and MCF10A, cells were plated at 30k cells / cm^2^, and assayed 48 hrs post seeding. For short term lentivirus transduction, cells were plated at 20k cells / cm2, transduced with lentivirus next morning, and assayed 72 hrs post seeding. All cultures were grown in 300 μL media. For H1 hESCs, wells were coated with 300 μL of Matrigel coating solution (see hESC cell culture methods) for at least 2 hrs at 37 °C in tissue culture incubator, followed by seeding cells in 300 μL mTESR Plus media. Media was changed daily for hESCs.

For YAP-degron experiments, dTAGv1 (Tocris Cat. No. 6914) or DMSO control was added at 2 μM for various times (15, 30, 60min) in fresh media, followed immediately by EU incorporation.

EU incorporation assay in all conditions was performed by addition of 1 mM 5-Ethynyluridine (Fisher Scientific 50-238-7038) directly to wells, and incubated for 1 hr in tissue culture incubator. Following incorporation, samples were gently washed twice in PBS, fixed with 4% paraformaldehyde for 10 minutes at room temperature, washed 2 × 5 minutes in PBS, permeabilized for 5 minutes in 0.5% Triton X-100 in PBS, and washed 2 x 5 minutes in PBS.

The EU labeling reaction was carried out following the protocol in Jao & Salic ^10^. The labeling reaction buffer was made fresh from 100 mM Tris pH 8.0, 1 mM CuSO4, μM iFluor 488 Azide (AAT Bioquest Cat. No. 1000), and 100 mM ascorbic acid added last from 500 mM stock in water. 300 μL of reaction buffer was then added to 96-wells for 90 minutes at room temperature in the dark. 3 x 10 minute PBS washes were performed to thoroughly destain the unreacted azide dye. Cell nuclei were detected with DAPI (AAT Bioquest Cat. No. 17507) staining at 1 μM for 15 minutes.

### Microscopy segmentation and quantification

3D segmentation of cell nuclei, nucleoli, and sub-nuclear HSat3 foci was performed using custom image segmentation software (https://github.com/jmfrank/track_analyzer).

#### Nuclear segmentation

A 9X9X3 pixel median filter was performed, followed by intensity thresholding, hole-filling, watershedding, and object size filtering to identify distinct nuclei (all operations performed in 3D).

#### Nucleoli segmentation

The nucleoli were segmented by first using the nuclear segmentations as a positive mask, followed by a 3X3X1 median filter and a local intensity thresholding scheme. Nucleoli size and EU intensity is too variable between individual cells. Thus, the intensity threshold was determined on a per-cell basis. The nucleoli intensity threshold was set at 6 signal-to-noise ratio using the non-nucleolar pixels which empirically are the 10-60 intensity percentiles per cell nucleus. Object sizes are then filtered to obtain nucleoli masks. The nucleoli masks were used to measure mean intensity per nucleolus after subtracting a local background per nucleolus which is a 3D shell mask of width 2 pixels starting at 6 pixels from the nucleolus boundary (ignoring pixels from other nucleoli in the same nucleus).

#### Colocalization

ZFsat3-megfp foci were first segmented to provide a HSat3 reference location in cell nuclei. This segmented ROI was then used to analyze intensity of TF-mscarlet in a second channel. Because HSat3 arrays are variable in size, we first generated scatter plots to correlate signal intensities. Second, we generated 1D intensity profiles for the two channels by mapping 2D intensity to polar coordinates centered at the foci centroid position. This showed strong correlation for most top HSat3 candidates. However, SOX1 and NKX2-2 only showed colocalization for a small set of foci, so 1D line-profiles were generated manually using Fiji.

Image processing scripts provided upon request.

### Total RNA / DNA ratio in TEAD knock-out cells

To estimate the impact of TEAD1-4 knock-out on total RNA expression, we estimated the total RNA to total DNA for cultures of MCF7 and MCF7 TEAD1-4 knockout (TEAD-qKO)^60^. Approximately 200,000-400,000 cells were collected in triplicate for each cell line using TrypLE Express. Cultures were split into equal volume and total RNA (Qiagen RNeasy Plus Universal Mini Kit Cat no. 73404) and total DNA (Qiagen Blood & Cell Culture DNA Mini Kit Cat no. 13323) were measured.

## DATA AVAILABILITY

Raw data from DiMeLo-seq and microscopy experiments are available upon request and will be deposited in permanent repositories prior to publication. The library of mScarlet-tagged TFs will be deposited in Addgene prior to publication.

## CODE AVAILABILITY

Custom scripts for performing data analysis for the TF motif and ChIP-seq screens will be made publicly available on github and in permanent repositories upon publication.

## ACKNOWLEDGEMENTS

We thank Nathan Gamarra for the many fruitful discussions for analyzing and interpreting data, as well as Sylvia Erhardt and Samuel Corless for useful discussions regarding ChIP-seq analysis. *Core facilities:* Stowers Institute Microscopy Core, Stanford University Cell Sciences Imaging Core Facility (RRID:SCR_017787). Figure 3a was created in BioRender, https://BioRender.com/s4wcmqh.

## CONTRIBUTIONS

JMF conceived the project. JMF, DD, NA, KLG, JLG designed the computational and experimental strategies. JMF performed the motif search and analysis. DD performed the ENCODE ChIP-seq analysis. JMF generated TF fusion constructs, microscopy data, stable expression cell lines, gene edits, and western blots. JMF performed image quantification. CC, AB, RJL, JMF performed DiMeLo-seq, ONT sequencing, and basecalling. JMF performed DiMeLo-seq analysis. RPG designed CRISPR knock-in strategy. JMF and NA wrote and edited the manuscript. JMF and NA generated figures. KLG, NA, and JLG provided project funding and supervision.

## SUPPLEMENTARY FIGURES

**Supplementary Figure 1:**
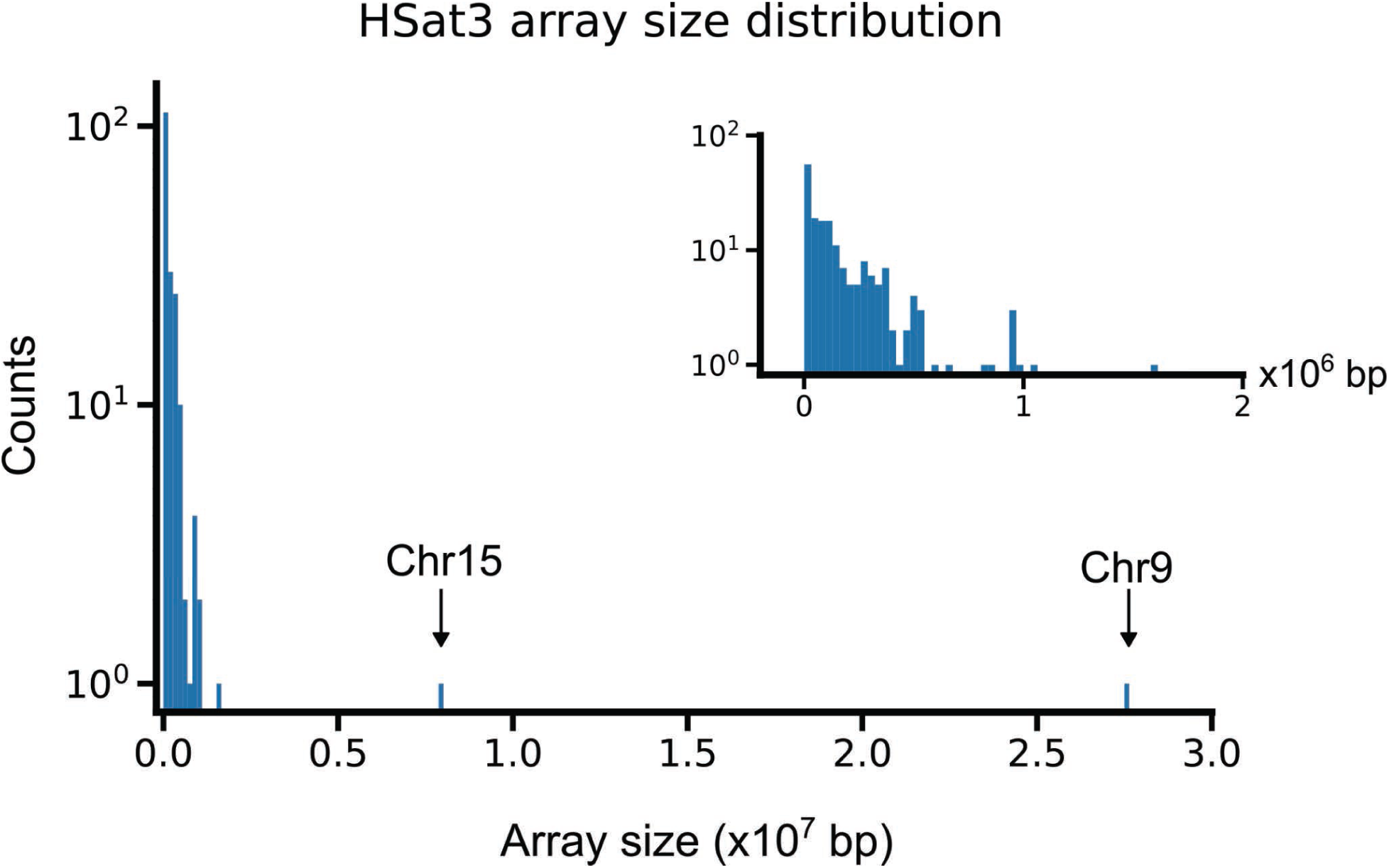
Distribution of HSat3 array sizes in T2T-CHM13v2.0.

**Supplementary Figure 2:**
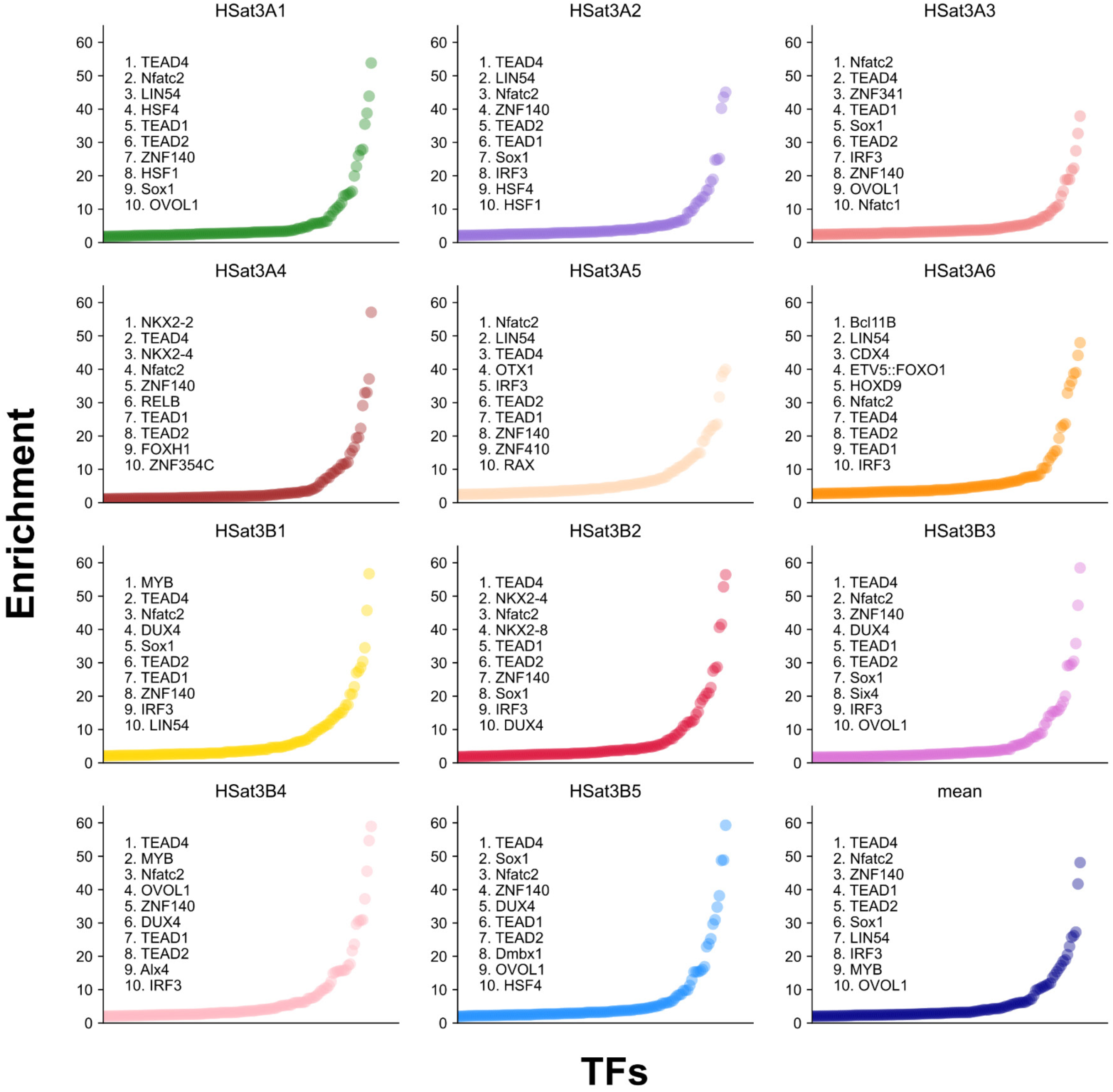
Ranked HSat3 motifs for all sub-families.

**Supplementary Figure 3:**
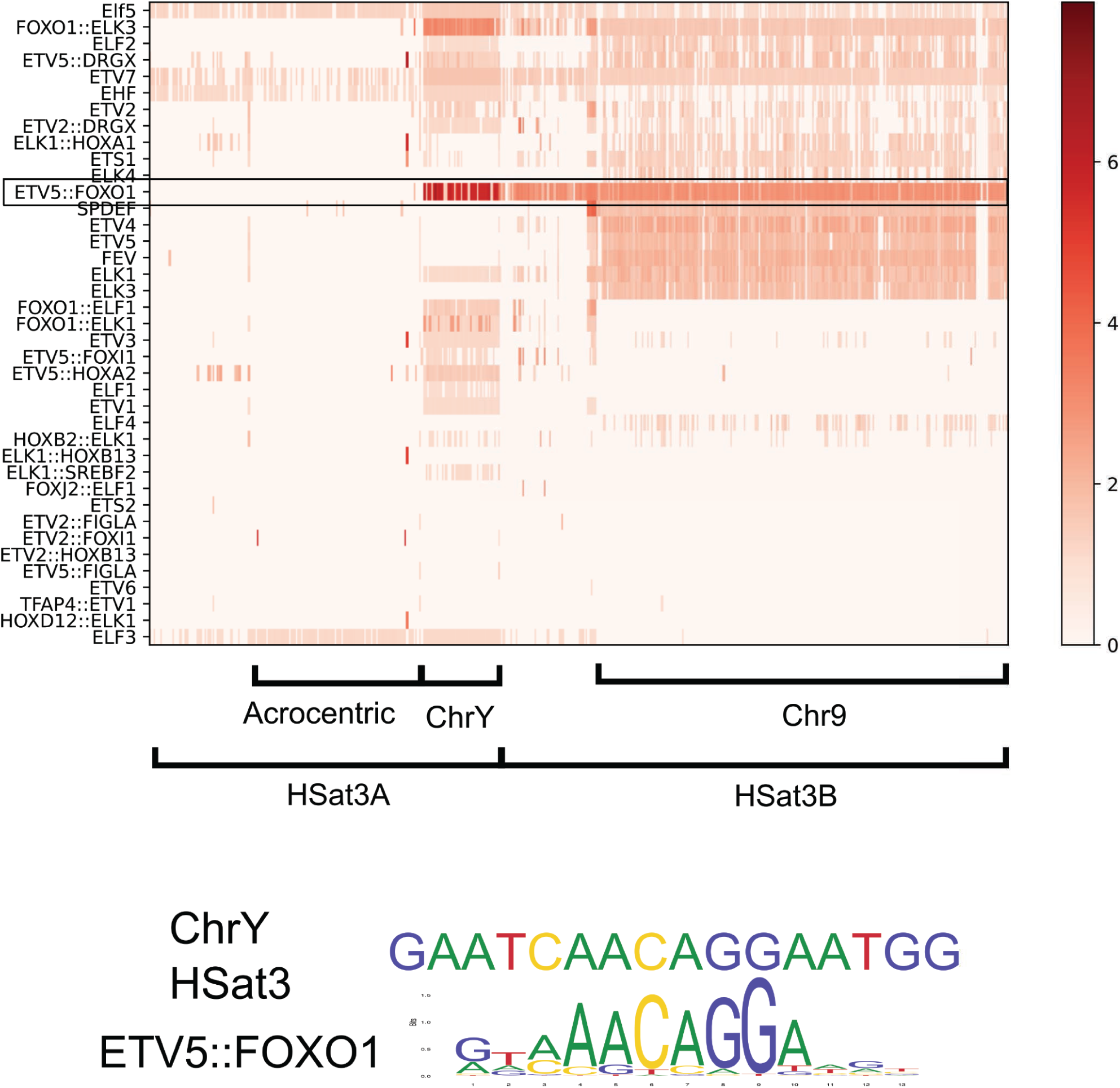
ETS-family motifs in HSat3.

**Supplementary Figure 4:**
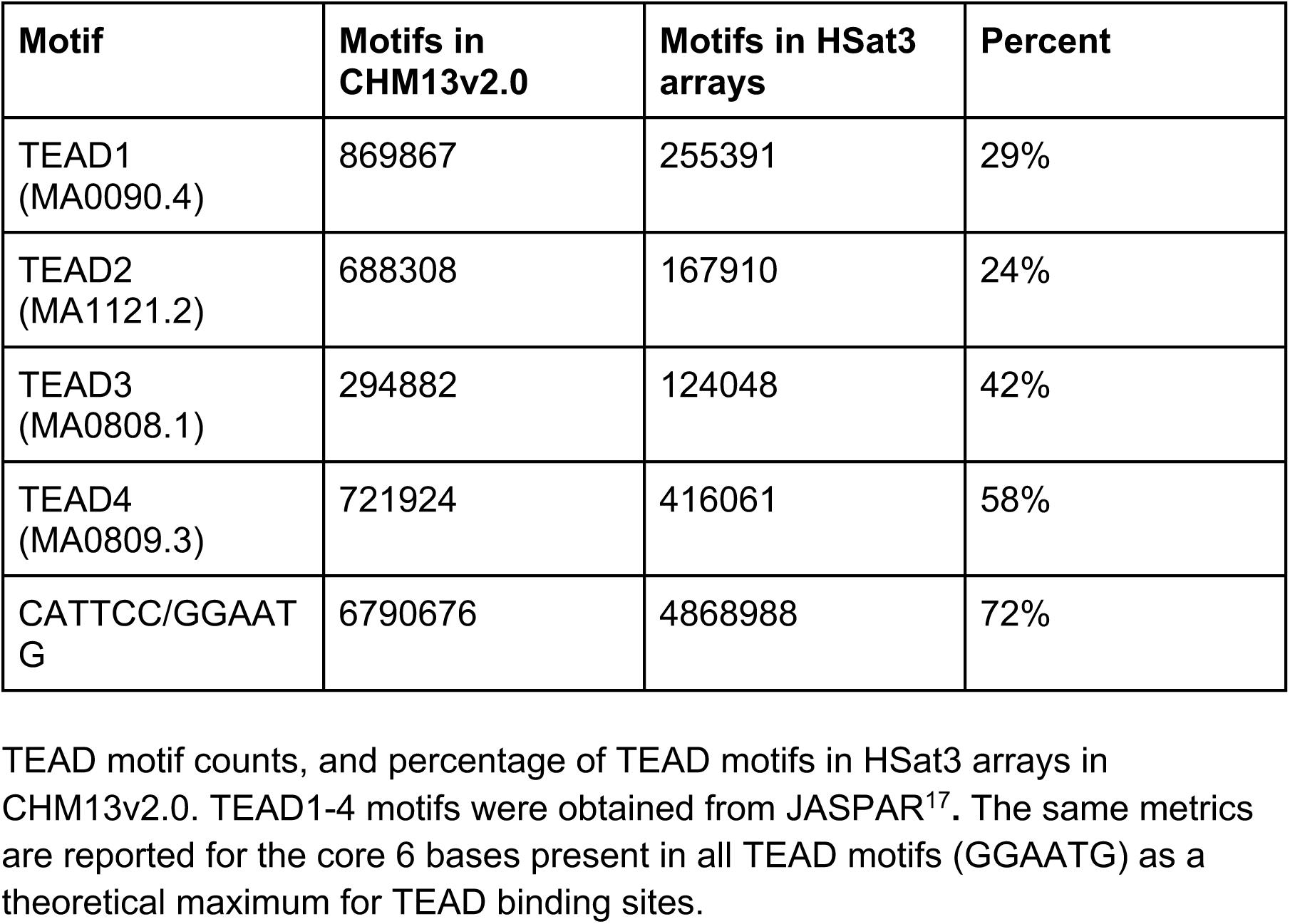
Estimation of TEAD family motifs within HSat3.

**Supplementary Figure 5:**
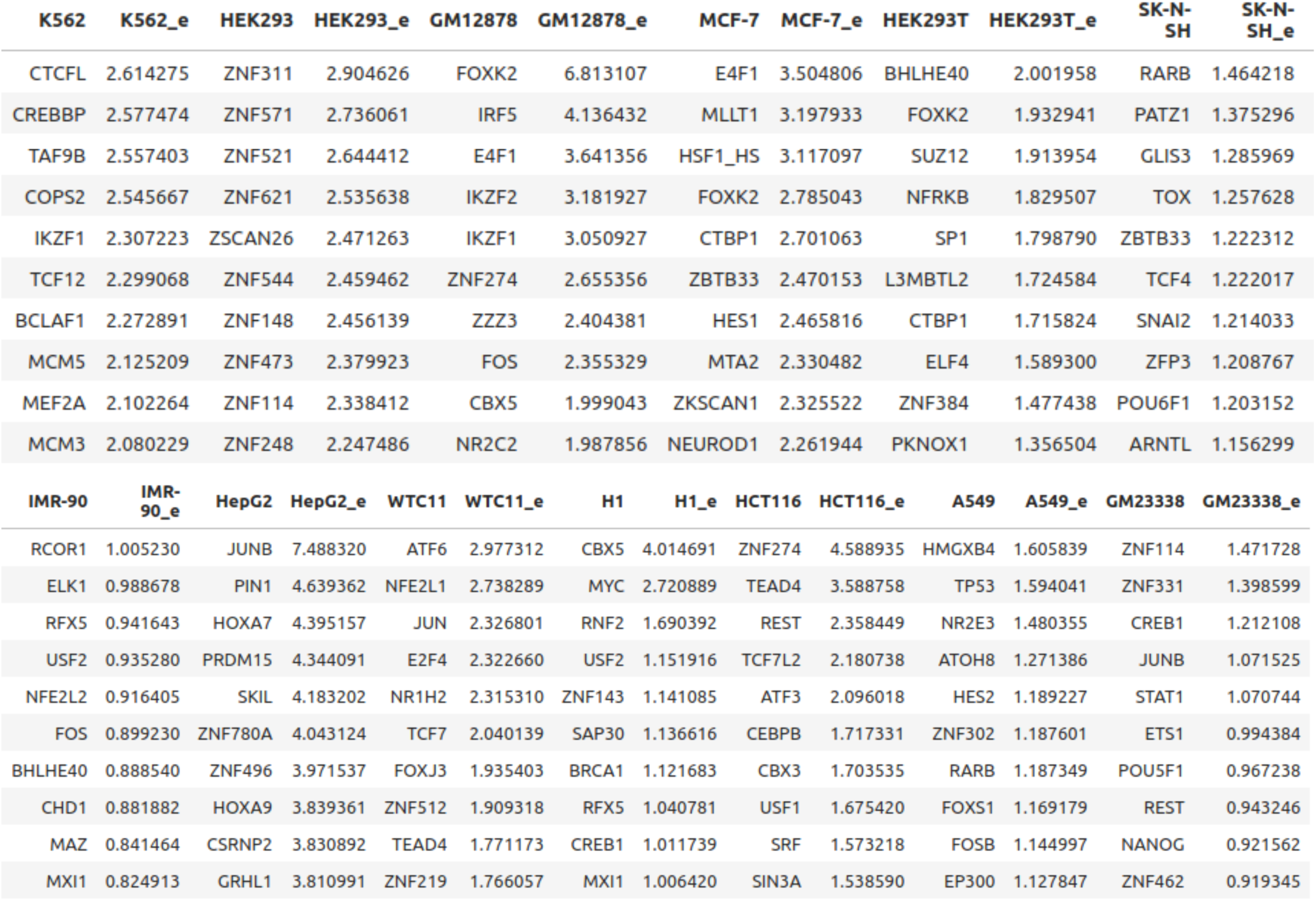
Top 10 HSat3-enriched factors per cell line using ENCODE ChIP-seq data from 13 cell lines. Columns are pairs of cell line followed by the enrichment score. Note, not all factors are tested in all cell lines. Full dataset can be found in supplementary data.

**Supplementary Figure 6:**
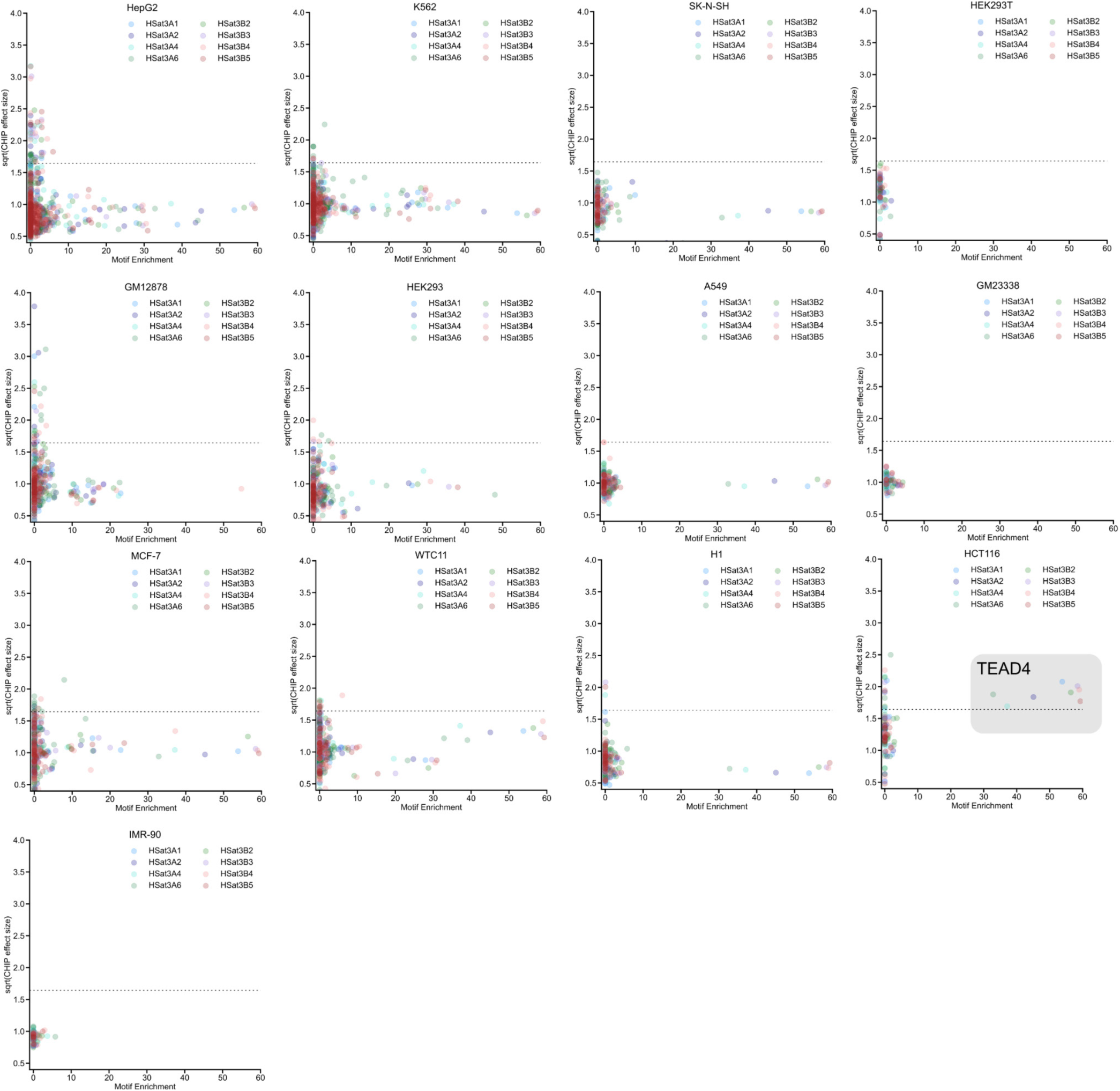
Correlation between ChIP-seq and motif enrichment analysis. Data points are colored by the HSat3 subfamily, and represent the mean value of a given factor’s enrichment in ChIP-seq and motif analysis, for each cell line. The dashed line represents an enrichment cut-off threshold generated by HSF1 data in heat-shocked MCF7 cells (Figure 2b-c).

**Supplementary Figure 7:**
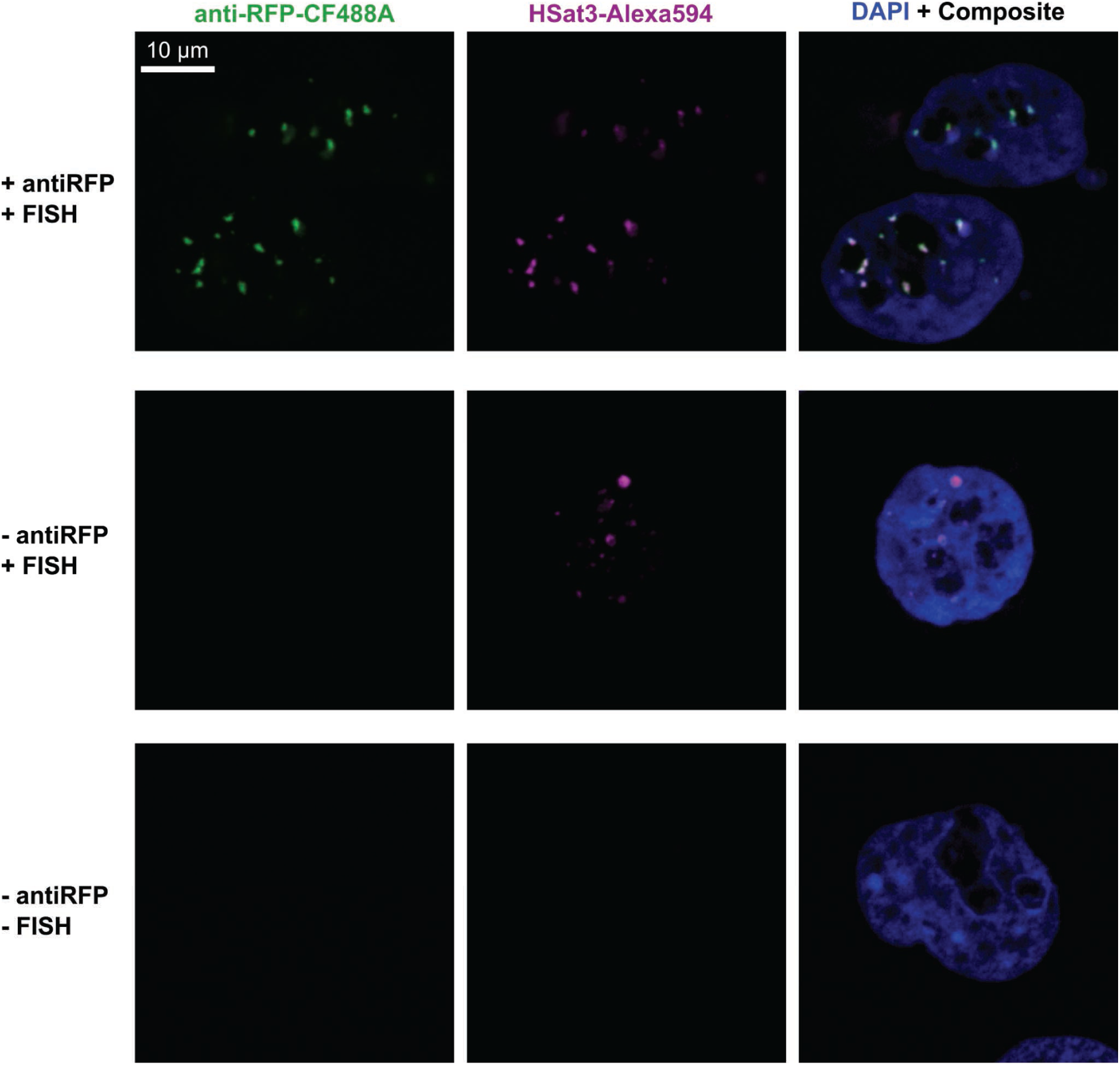
Immuno-FISH Validation of ZFsat3-mScarlet using HSat3-FISH. MCF10A cells stably expressing ZFsat3-mScarlet were prepared for immuno-FISH with 488-labeled anti-RFP, Alexa594-labeled HSat3 oligos with two control samples: (1) HSat3 FISH only and (2) neither anti-RFP or HSat3 FISH.

**Supplementary Figure 8:**
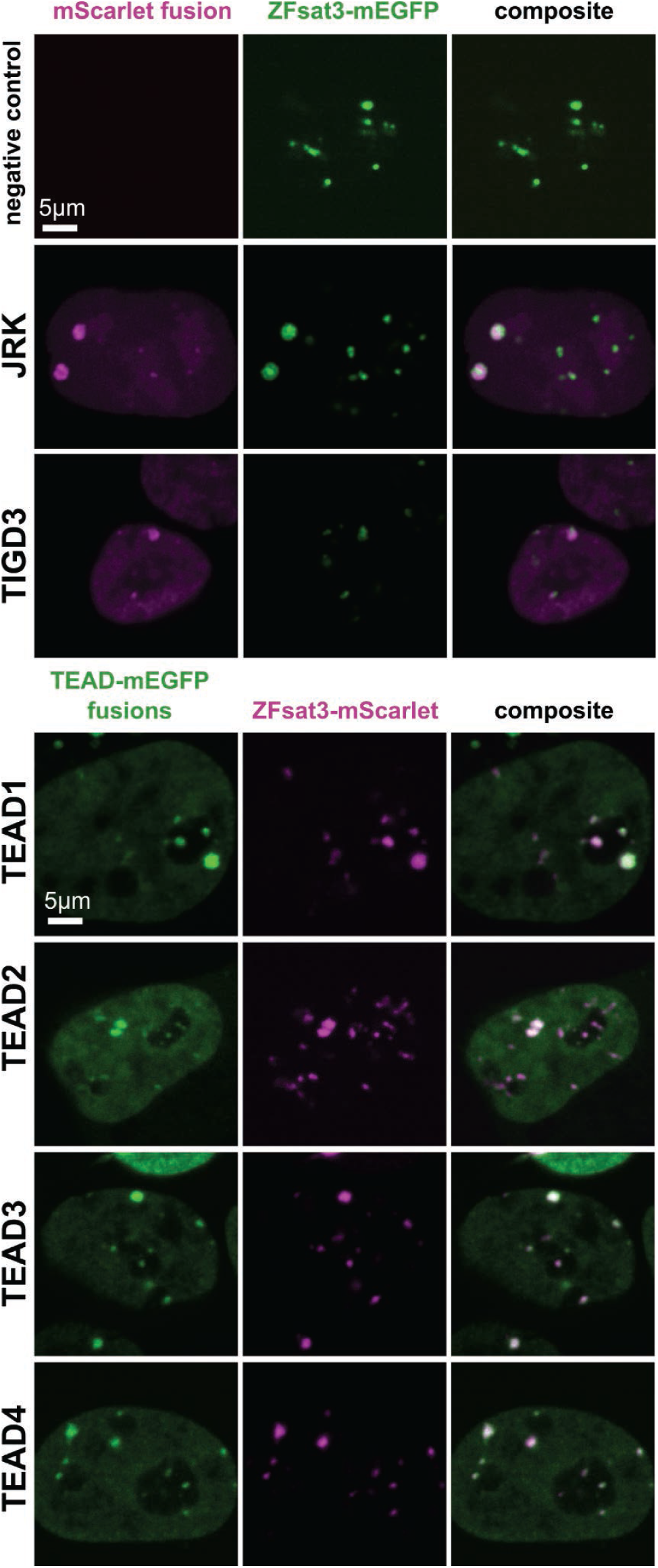
JRK, TIGD3, and TEAD-family factors colocalize with HSat3. Negative and positive controls for imaging screens. Cells transfected with ZFsat3-mEGFP do not show strong bleedthrough into mScarlet channel. JRK and TIGD3 are previously reported HSat3 binding factors, both showing colocalization with ZFsat3-mEGFP.

**Supplementary Figure 9:**
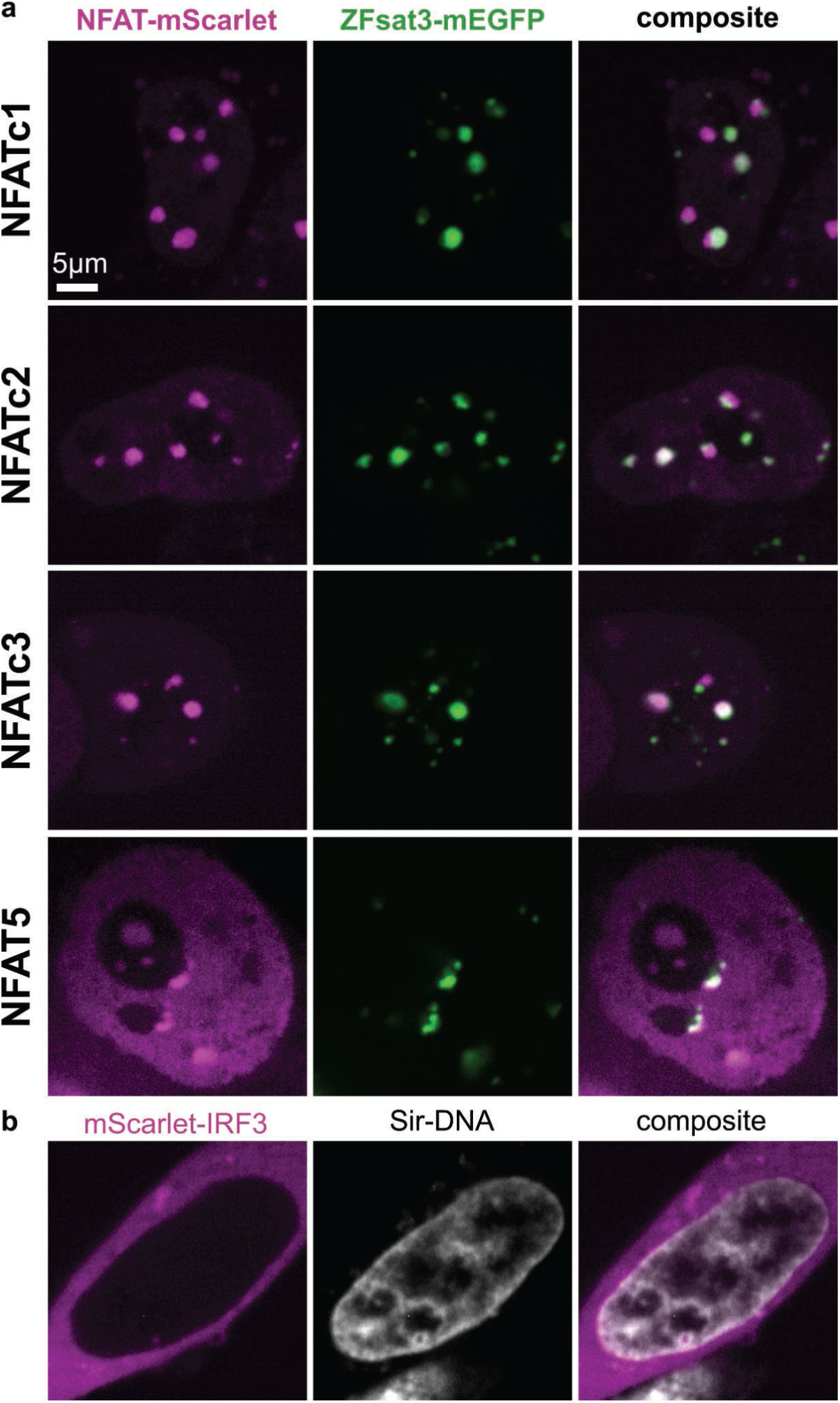
(a) Ionomycin-stimulated NFAT-family factors colocalize to ZFsat3. (b) mScarlet-IRF3 is cytoplasmic in transfected 293T cells.

**Supplementary Figure 10:**
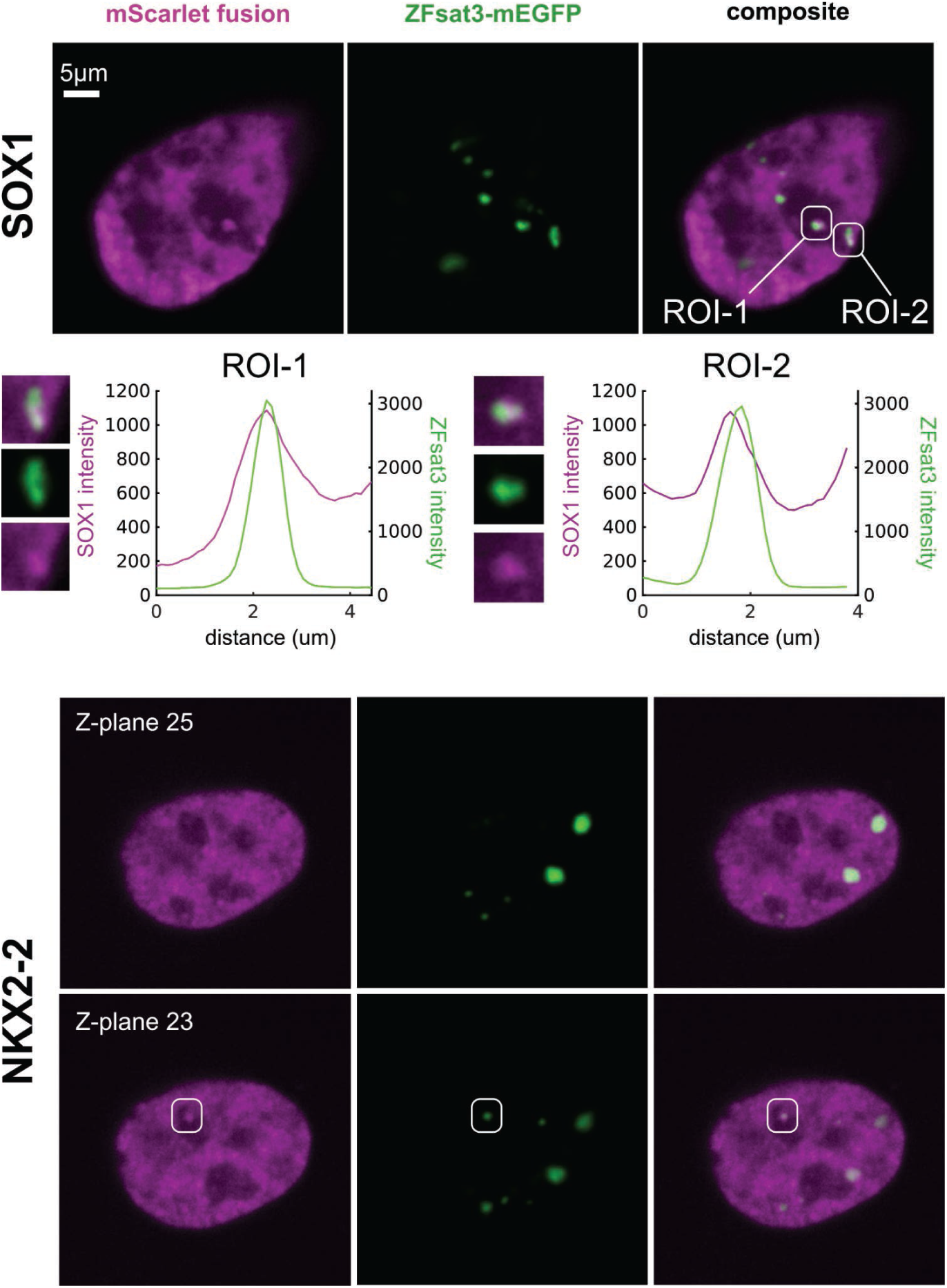
Subtle colocalization of SOX1 and NKX2-2 with ZFsat3-mEGFP. ROIs show a subset of foci are colocalized with HSat3.

**Supplementary Figure 11:**
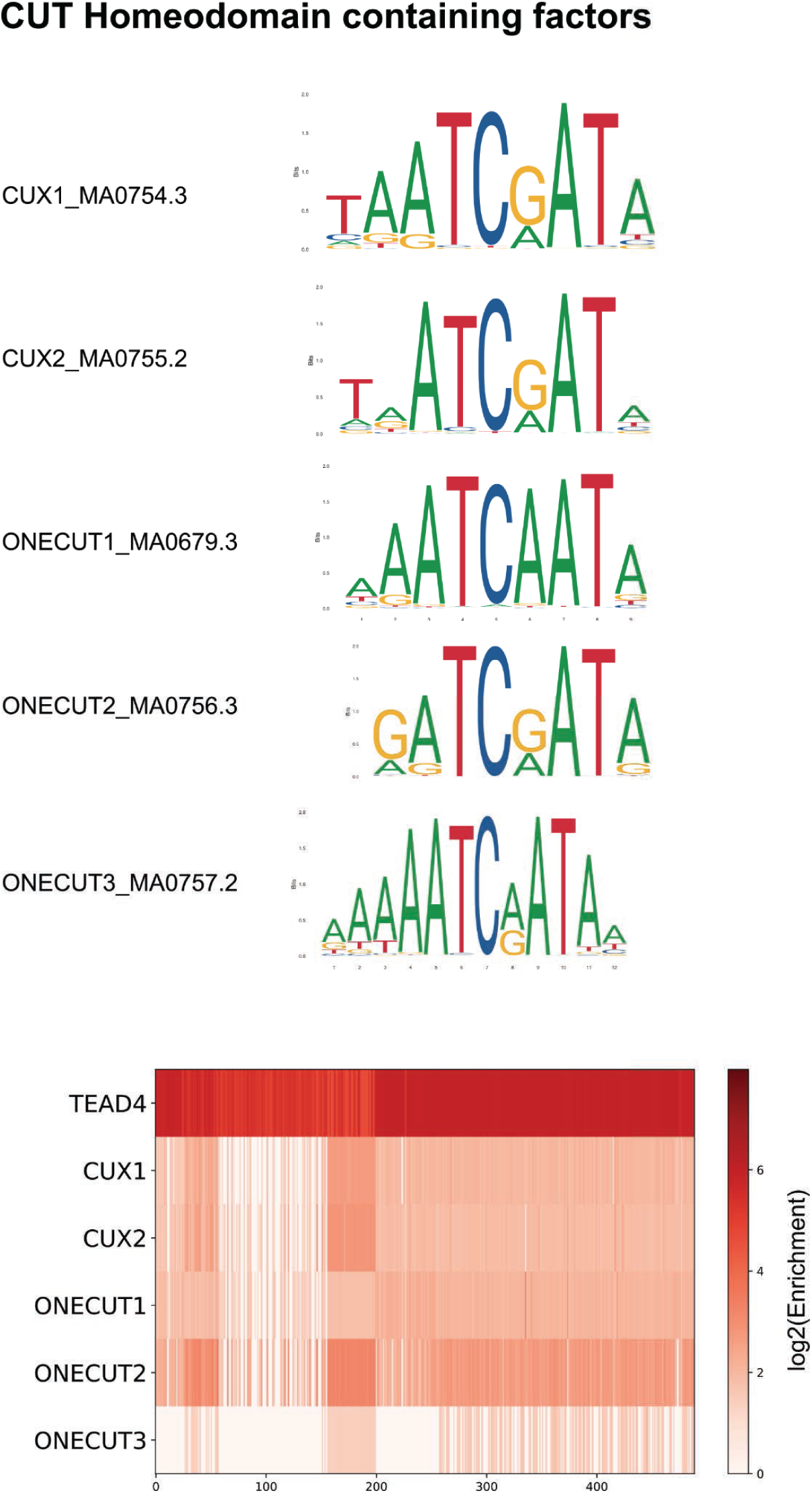
Motifs for CUT Homeodomain containing factors. TF motif enrichment shows weak enrichment of motifs in HSat3 compared to TEAD4, however, imaging showed strong HSat3-binding (Figure 3b).

**Supplementary Figure 12:**
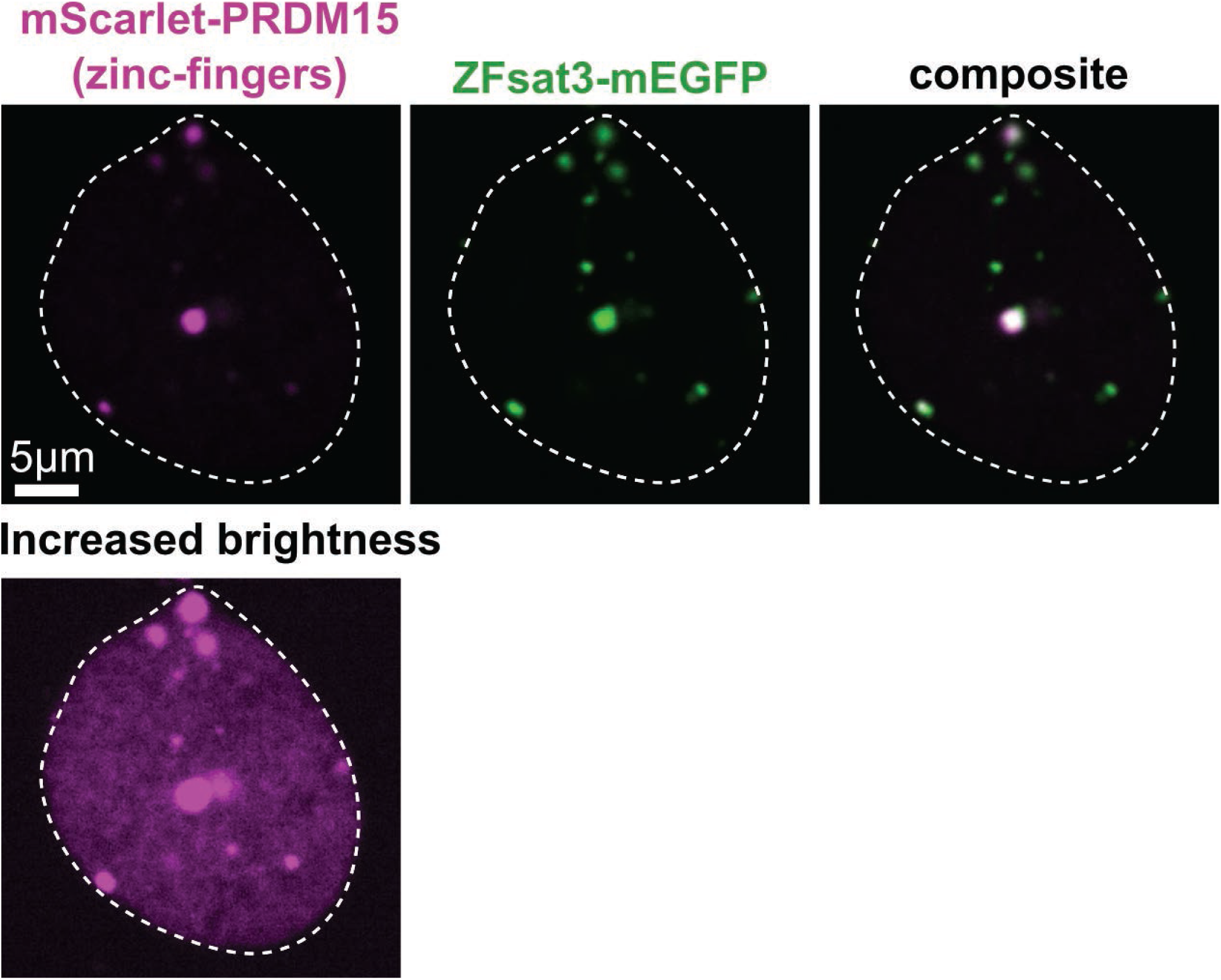
The PRDM15 ZF array binds HSat3. The nucleus is shown by increasing the display brightness.

**Supplementary Figure 13:**
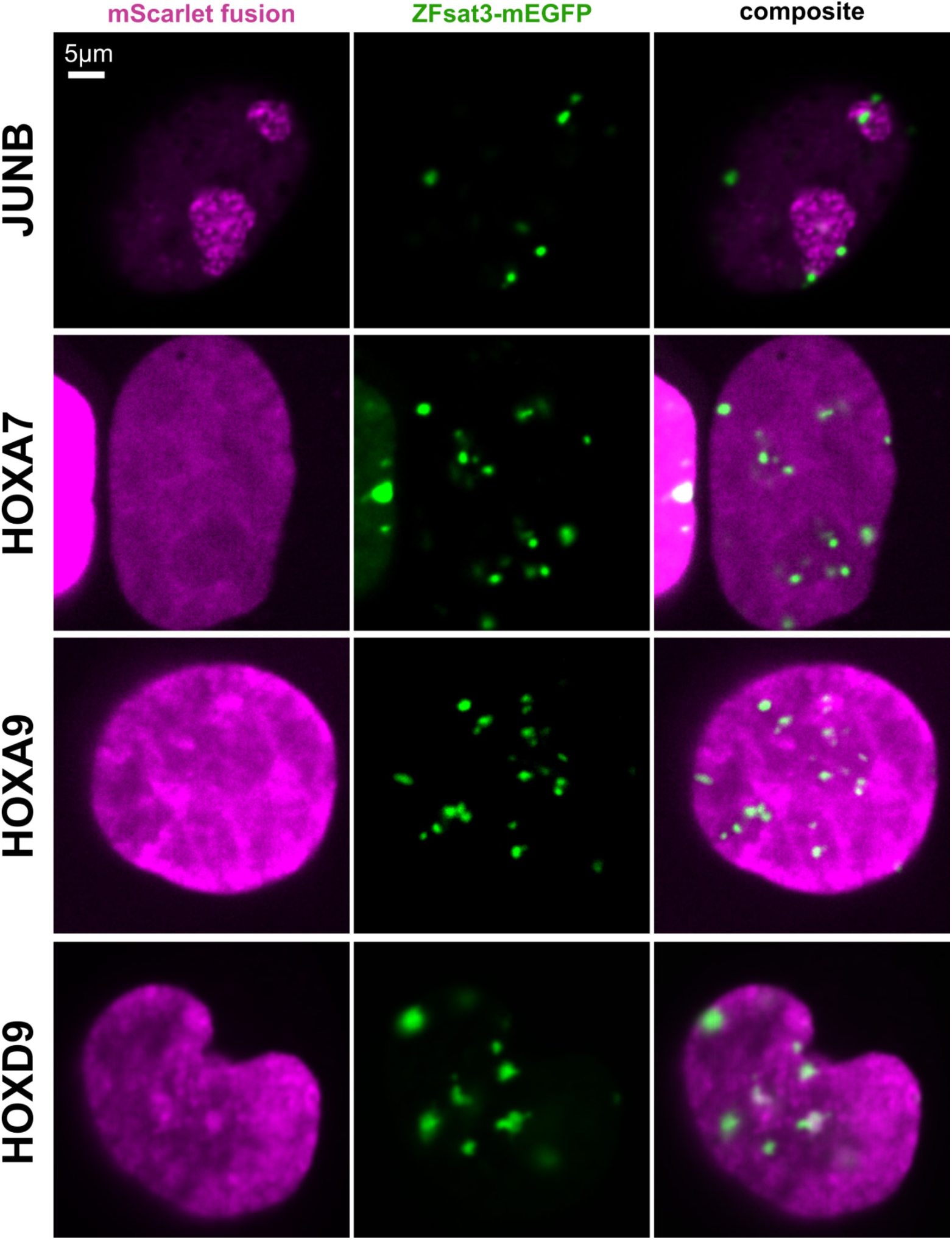
Colocalization of ChIP-seq HSat3 candidates JUNB, HOXA7, HOXA9, HOXD9 with ZFsat3.

**Supplementary Figure 14:**
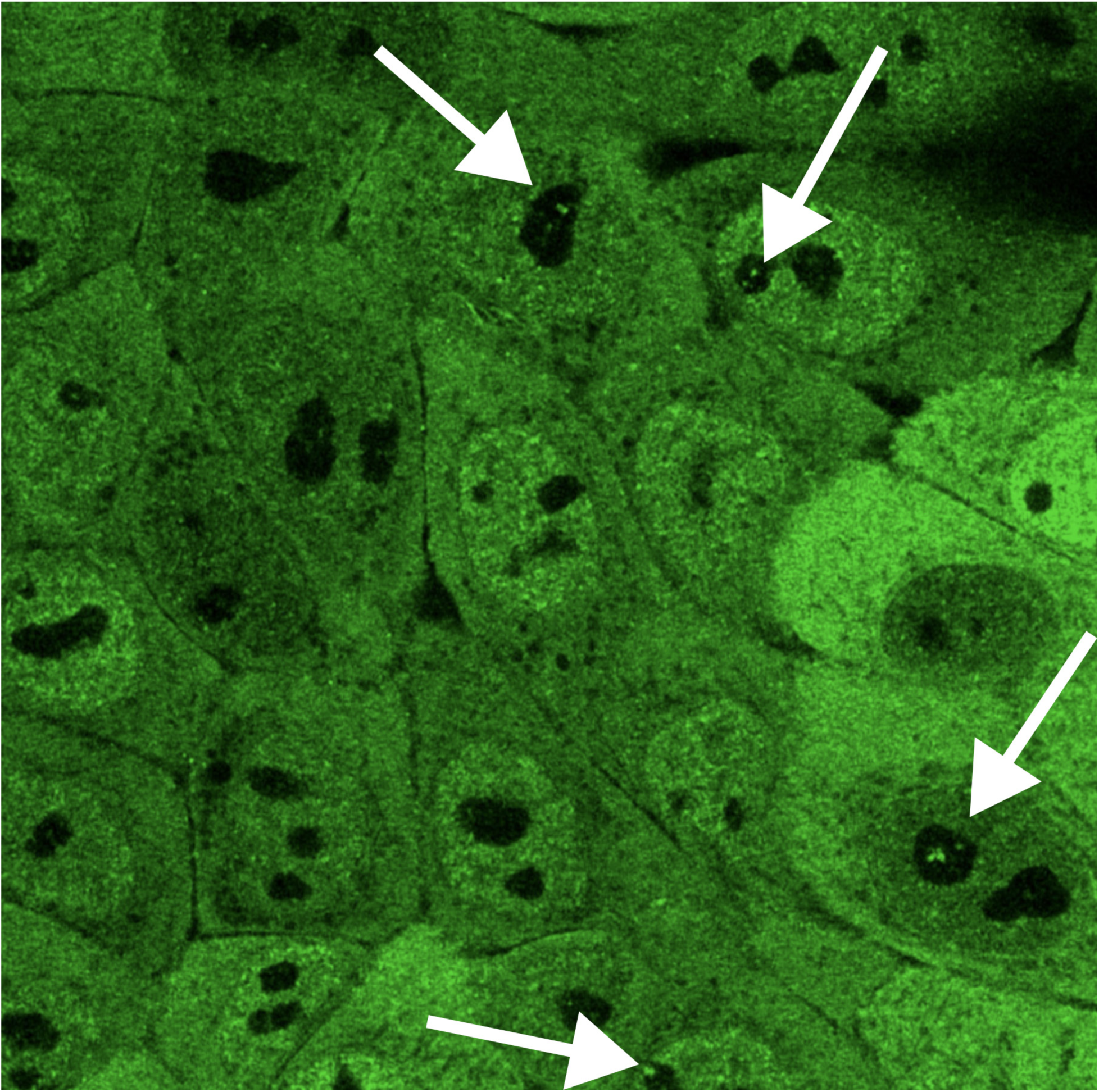
A single z-plane of live-cell super-resolution (Airyscan) microscopy of endogenous YAP-eGFP in MCF10A cells. Arrows point to bright, nucleolar YAP foci.

**Supplementary Figure 15:**
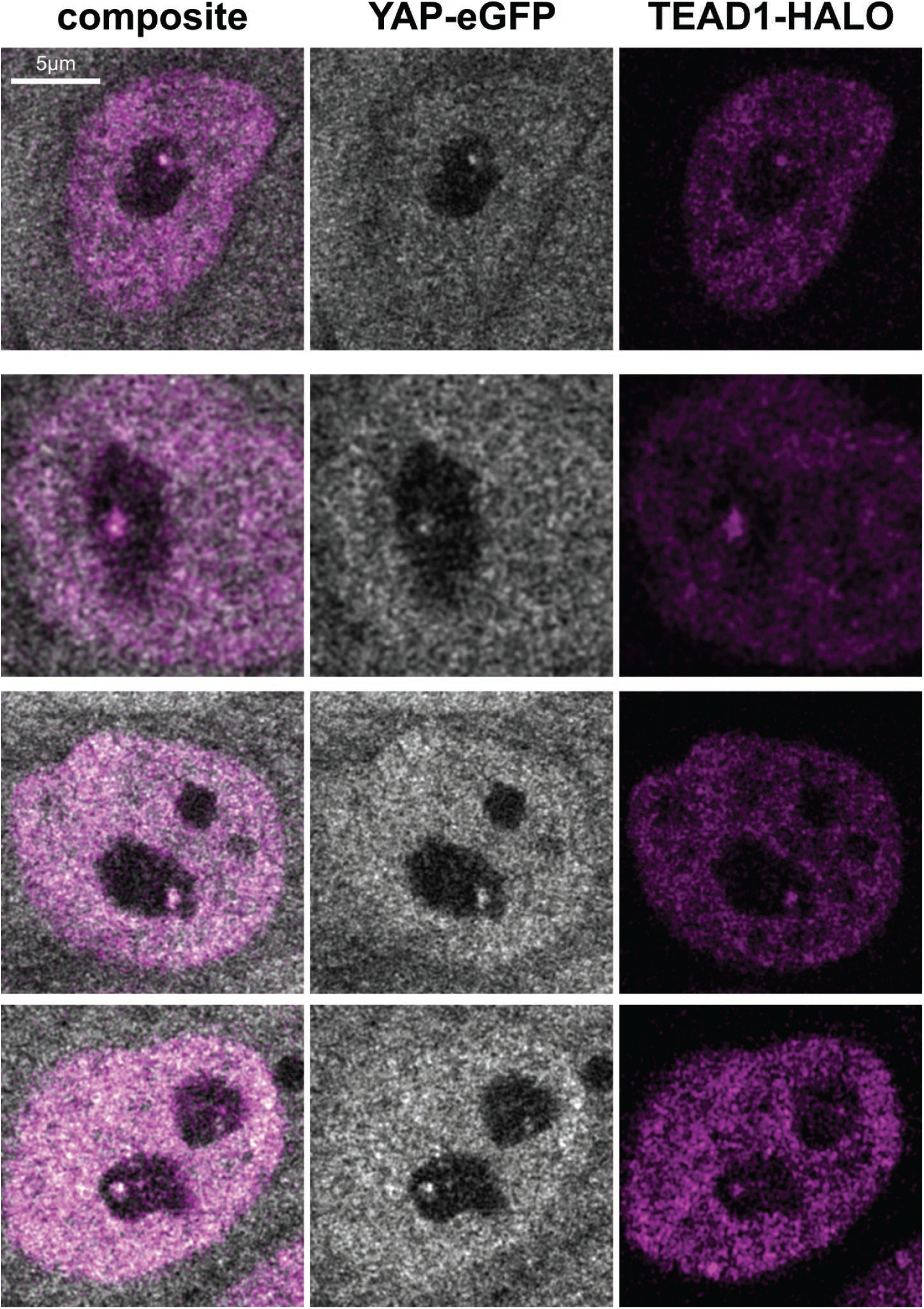
Colocalization of endogenous, nucleolar YAP-eGFP + TEAD1-HALO foci in H1 hESCs using Airyscan live-cell super-resolution imaging.

**Supplementary Figure 16:**
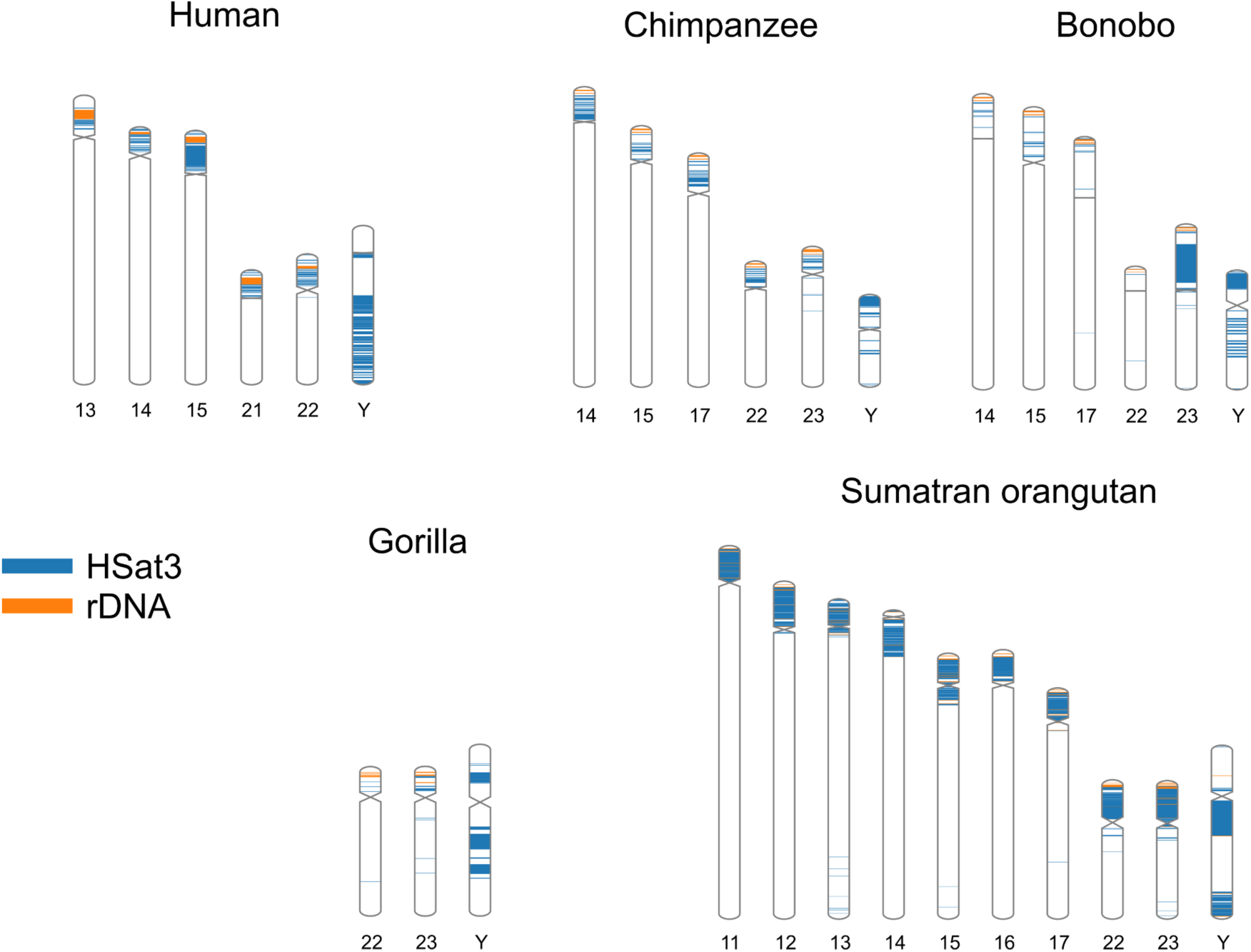
Organization HSat3 and rDNA arrays across great apes. For each species, the chromosomes containing rDNA are depicted, with colored blocks for HSat3 and rDNA. Chromosome Y is included for all species. Chromosome 13 in Orangutans contains both HSat3 and rDNA. Whereas the human homolog, chromosome 9, only contains HSat3 (Figure 1a), suggesting the human ancestral state contained rDNA.

**Supplementary Figure 17:**
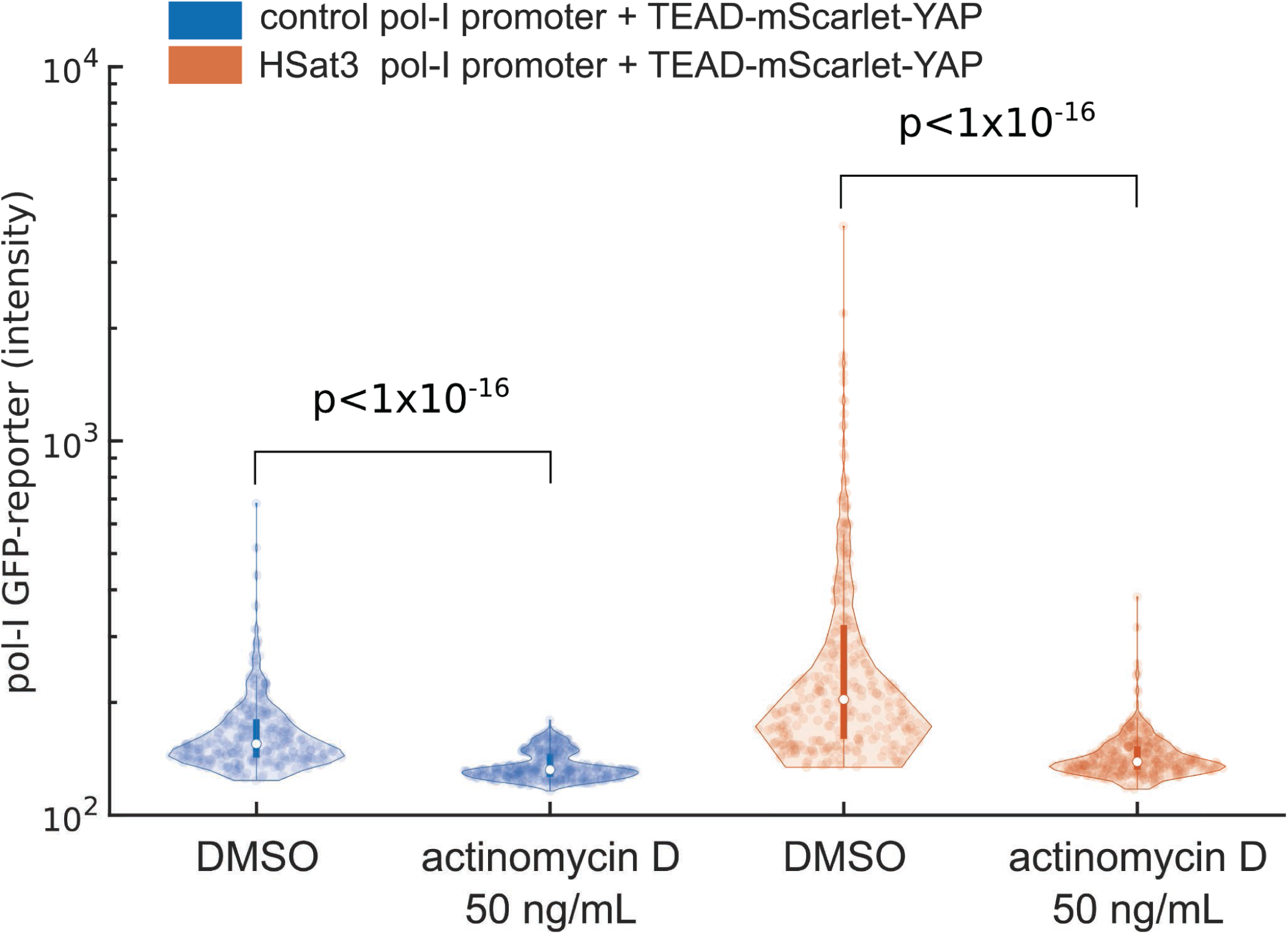
rDNA-pol-I reporter assays are sensitive to actinomycin D at 50 ng/mL. Data points represent individual cells. The number of cells per condition are as follows: N(control DMSO)=314, N(control actD)=400, N(HSat3 DMSO)=314, N(HSat3 actD)=410. Data is from one biological replicate. Statistical significance was assessed using a one-tailed Wilcoxon rank sum test. Experiment was repeated twice with similar results.

**Supplementary Figure 18:**
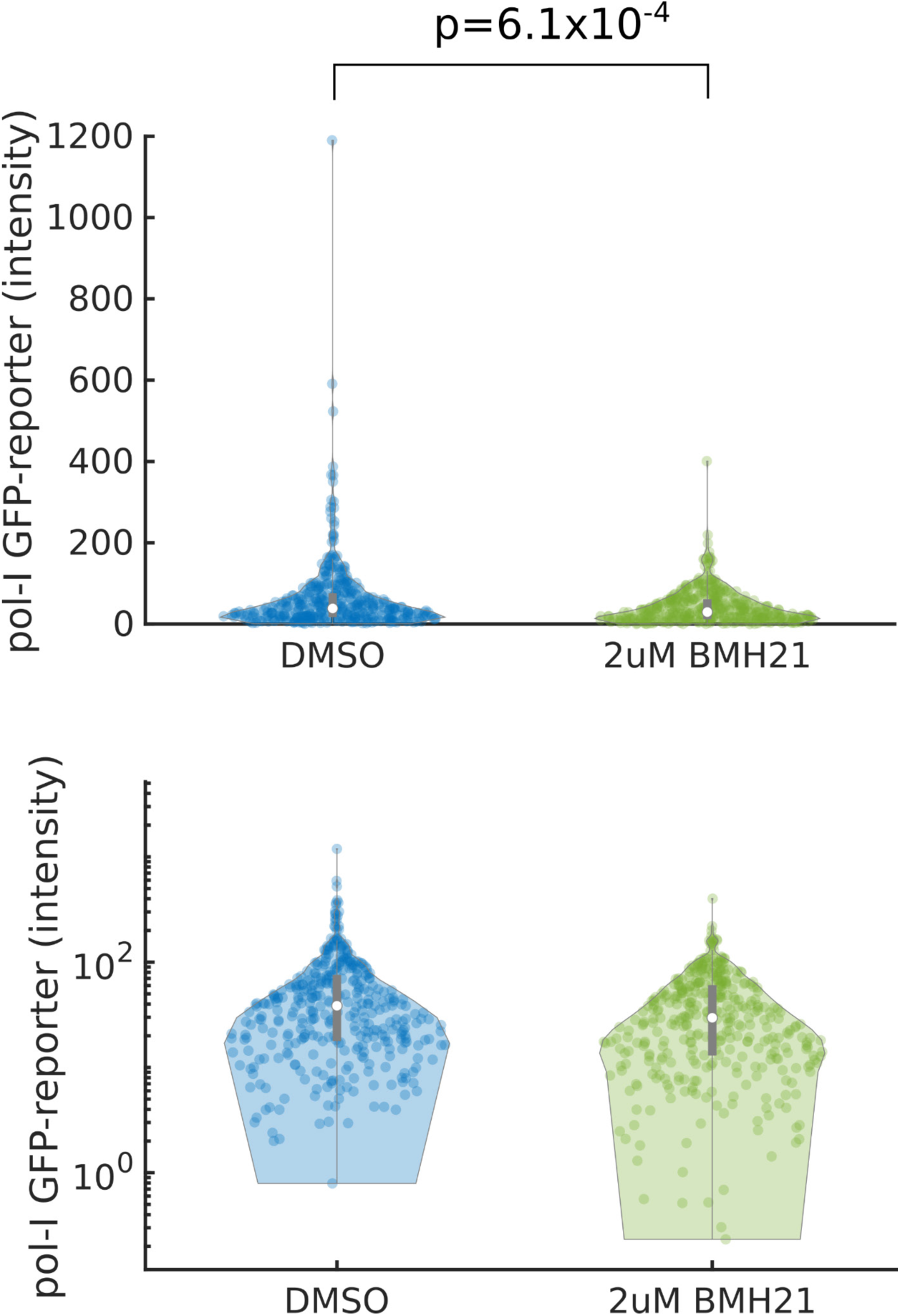
rRNA pol-I reporter assay is sensitive to Bmh21 at 2 μM. Data points represent individual cells. Following background correction due to autofluorescence of BMH21, 376 segmented cells remained with above zero signal intensity. We thus randomly selected 376 cells from the DMSO condition to compare reporter activity. Data is from one biological replicate. Statistical significance was assessed using a one-tailed Wilcoxon rank sum test. Experiment was repeated twice with similar results.

**Supplementary Figure 19:**
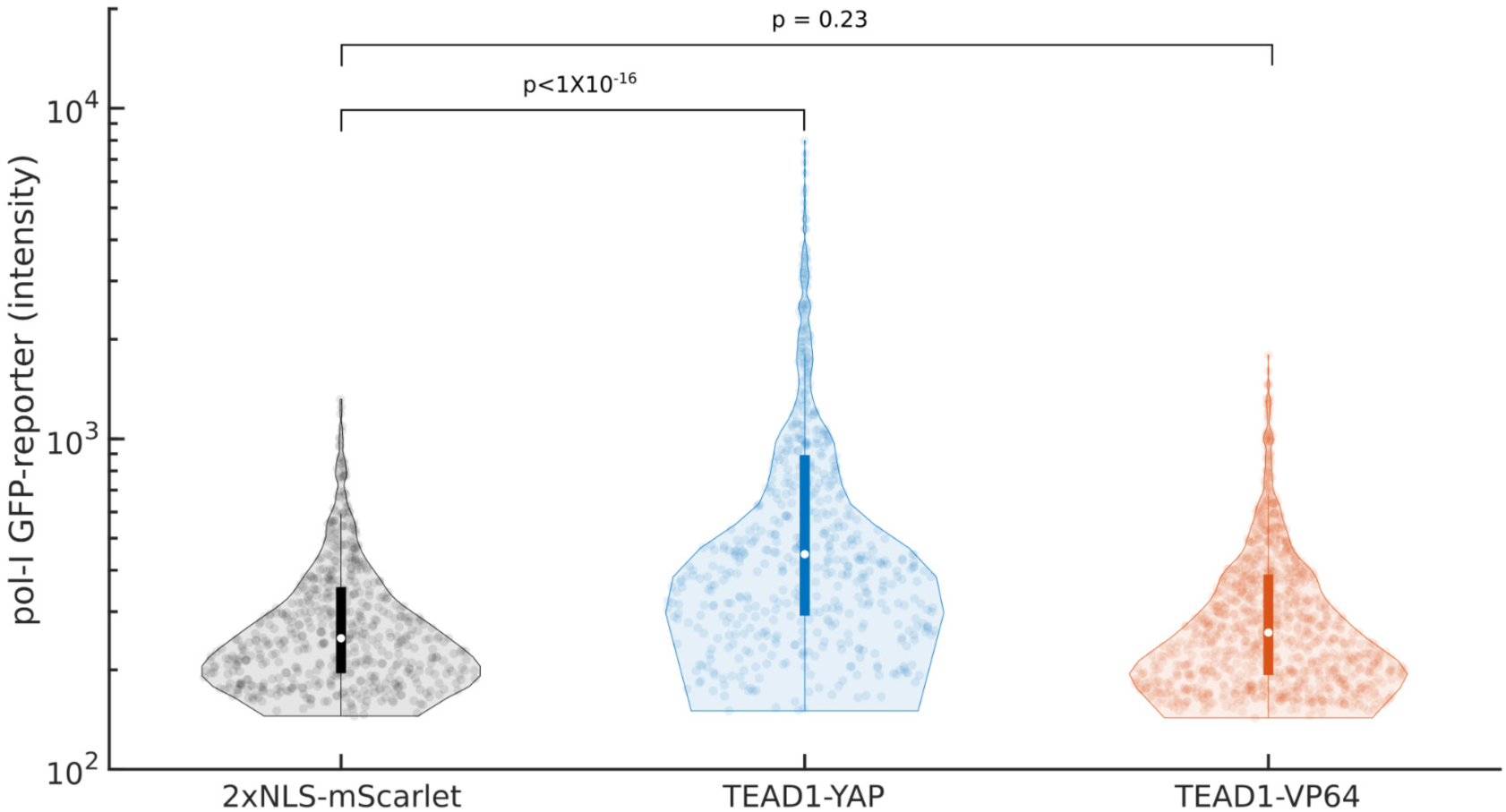
Activity of HSat3-rDNA enhancer when co-transfected with 2xNLS-mScarlet, TEAD1-YAP, and TEAD1-VP64. Data points represent individual cells. The number of cells per condition are as follows: N(NLS-mscarlet)=546, N(TEAD1-YAP)=556, N(TEAD1-VP64)=993. Data is from one biological replicate. Statistical significance was assessed using a one-tailed Wilcoxon rank sum test. Experiment was repeated twice with similar results.

**Supplementary Figure 20:**
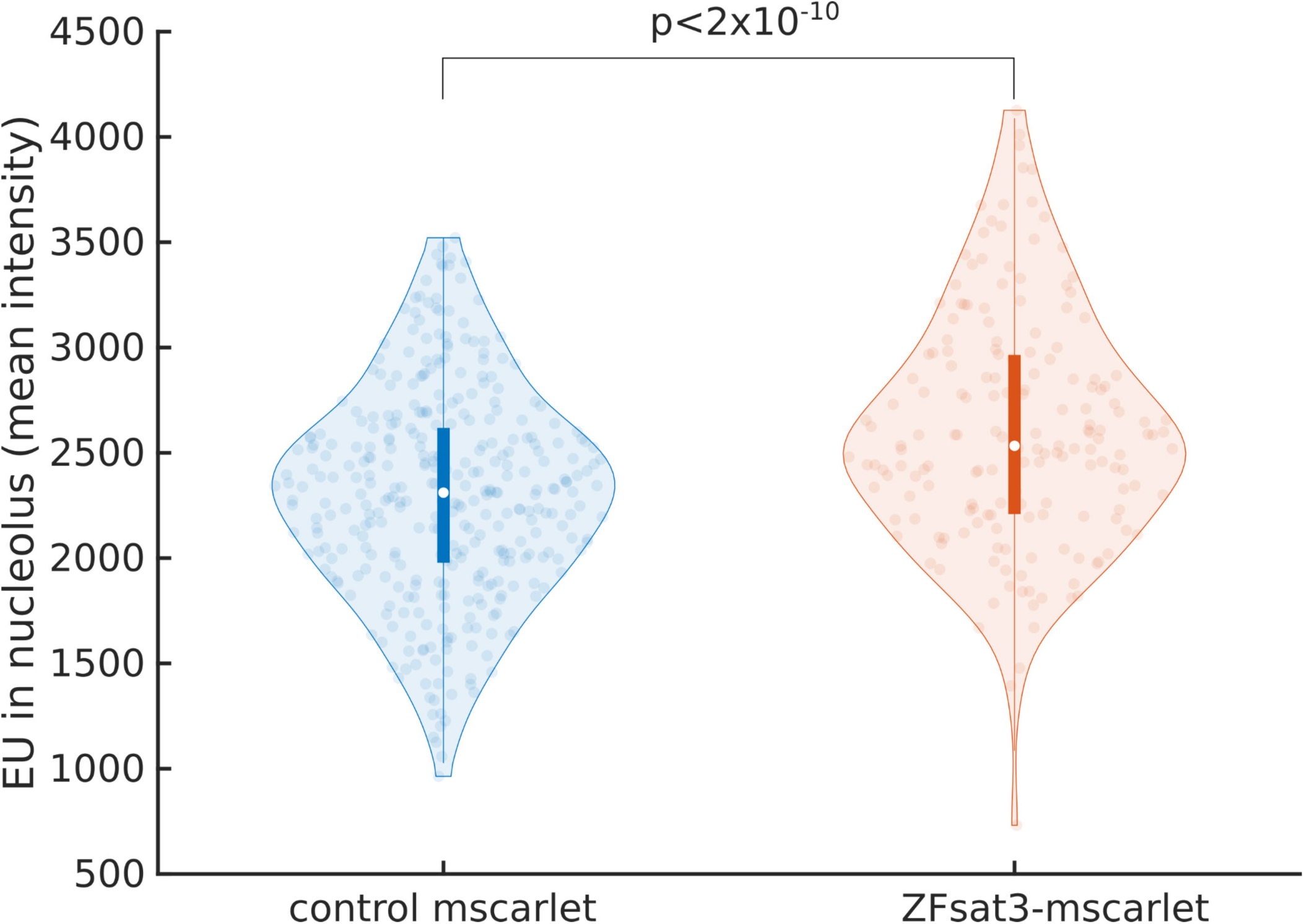
ZFsat3-mScarlet does not impact rRNA transcription in MCF10A cells. Expression of ZFsat3 only mildly increases EU signal, whereas ZFsat3-KM strongly represses EU signal (Figure 6d-e). Data points represent intensity values of segmented nucleoli. The number of nucleoli per condition are as follows: N(control mscarlet)=349, N(ZFsat3-mscarlet)=175. Data is from one biological replicate, repeated twice with similar results.

**Supplementary Figure 21:**
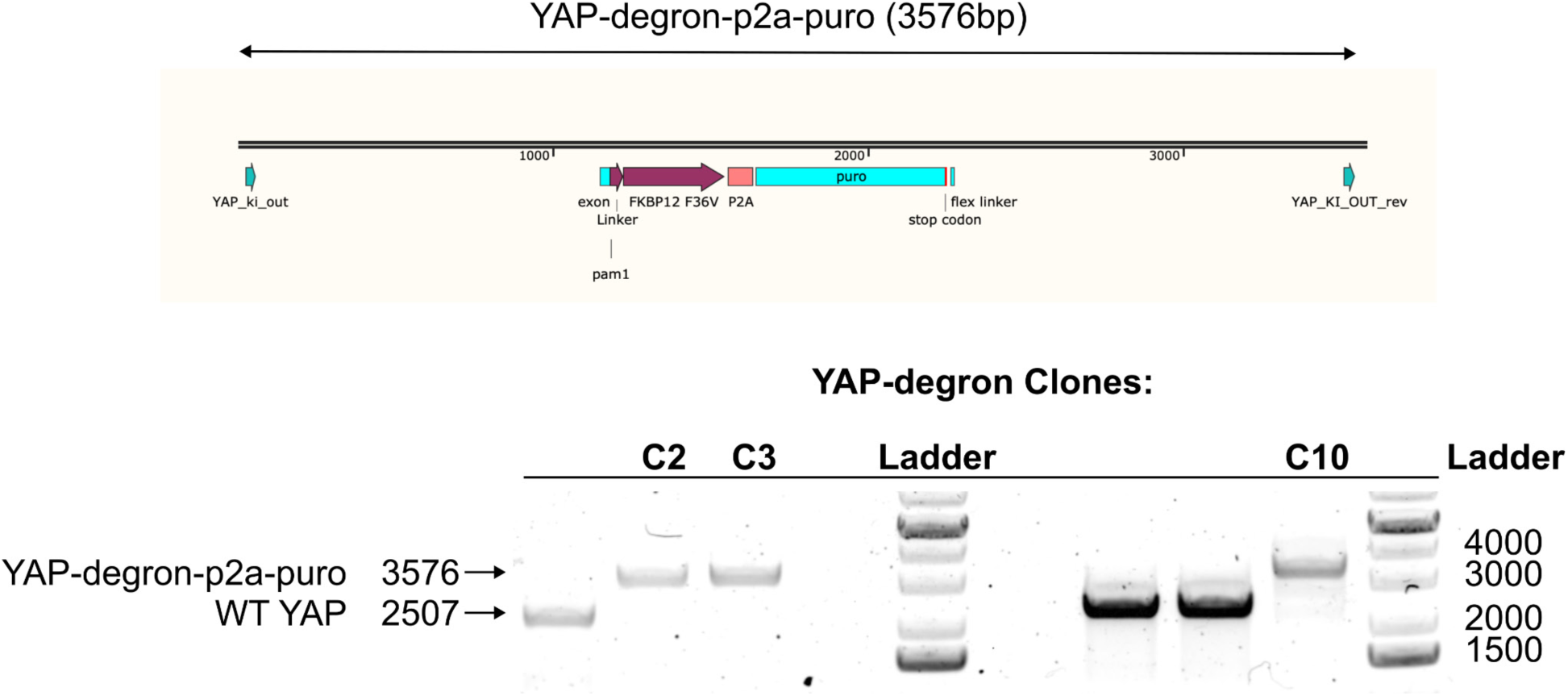
Genomic PCR validation of YAP-degron clones in H1 hESC cells. Full length PCR amplicon for YAP-degron-p2a-puro is 3576 base pairs. Clones 2, 3, and 10 show a single band at the knock-in mark indicating homozygous knock-in.

## SUPPLEMENTARY DATA

**[1]** “motif_enrichments_HSat3.csv”: Data table of DNA binding motif enrichments for each HSat3 satellite element in the reference genome chm13v2.0 and 828 DNA-binding factors.

**[2]** “chip_seq_enrichments_HSat3.csv”: Data table of mean HSat3 enrichment from ChIP-seq datasets obtained from ENCODE and aligned to chm13v2.0.

